# Loops and the activity of loop extrusion factors constrain chromatin dynamics

**DOI:** 10.1101/2020.02.29.969683

**Authors:** Mary Lou P. Bailey, Ivan Surovtsev, Jessica F. Williams, Hao Yan, Tianyu Yuan, Kevin Li, Katherine Duseau, Simon G. J. Mochrie, Megan C. King

## Abstract

The chromosomes - DNA polymers and their binding proteins - are compacted into a spatially organized, yet dynamic, three-dimensional structure. Recent genome-wide chromatin conformation capture experiments reveal a hierarchical organization of the DNA structure that is imposed, at least in part, by looping interactions arising from the activity of loop extrusion factors. The dynamics of chromatin reflects the response of the polymer to a combination of thermal fluctuations and active processes. However, how chromosome structure and enzymes acting on chromatin together define its dynamics remains poorly understood. To gain insight into the structure-dynamics relationship of chromatin, we combine high-precision microscopy in living *Schizosaccharomyces pombe* cells with systematic genetic perturbations and Rouse-model polymer simulations. We first investigated how the activity of two loop extrusion factors, the cohesin and condensin complexes, influences chromatin dynamics. We observed that deactivating cohesin, or to a lesser extent condensin, increased chromatin mobility, suggesting that loop extrusion constrains rather than agitates chromatin motion. Our corresponding simulations reveal that the introduction of loops is sufficient to explain the constraining activity of loop extrusion factors, highlighting that the conformation adopted by the polymer plays a key role in defining its dynamics. Moreover, we find that the number loops or residence times of loop extrusion factors influences the dynamic behavior of the chromatin polymer. Last, we observe that the activity of the INO80 chromatin remodeler, but not the SWI/SNF or RSC complexes, is critical for ATP-dependent chromatin mobility in fission yeast. Taken together we suggest that thermal and INO80-dependent activities exert forces that drive chromatin fluctuations, which are constrained by the organization of the chromosome into loops.

## Introduction

Chromatin, which is comprised of DNA and its associated proteins, is confined within the cell nucleus of eukaryotes, where it takes on a compartmentalized, three-dimensional structure. While we understand the molecular details of nucleosomes, the fundamental unit of chromatin, only recently has a unified model for the intermediate scale of chromatin organization emerged, namely the existence of topologically associating domains (TADs). 100 kb to Mb scale TADs are now considered to be a characteristic unit of mesa-scale chromosomal folding, revealed largely through genome-wide chromosome conformation capture techniques (e.g., Hi-C) (Dekker, 2014; Dixon et al., 2016; Dixon et al., 2012; Lieberman-Aiden et al., 2009; Mizuguchi et al., 2014; Nora et al., 2012; Sexton et al., 2012).

The structure of TADs is thought to arise through characteristic chromatin looping interactions. The major molecular machinery responsible for chromatin loops is composed of members of the structural maintenance of chromosome (SMC) complex family of ATPases, most notably in the form of the cohesin and condensin complexes (Banigan and Mirny, 2020). Indeed, high levels of cohesin and condensin reside at boundaries between TADs at steady-state (Mizuguchi et al., 2014; Nora et al., 2012). Given the established ability of SMC complexes to topologically engage DNA strands (Haering et al., 2008; Terakawa et al., 2017), it has been widely suggested that SMC complexes are loop extrusion factors (LEFs) that drive the formation of chromatin loops, an activity that can be observed *in vitro* or in extracts (Davidson et al., 2019; Ganji et al., 2018; Golfier et al., 2020; Kim et al., 2019). Moreover, loss of functional cohesin in fission yeast leads to a complete loss of TADs as assessed by Hi-C through a mechanism distinct from its role in sister chromatid cohesion (Mizuguchi et al., 2014) while complete depletion of cohesin similarly leads to the loss of TADs in mammalian cells (Rao et al., 2017). LEF simulations further demonstrate that the dynamic extrusion of loops by SMC complexes combined with the existence of boundary elements encoded in the DNA (for example, sequence-specific binding of CTCF, which is enriched at boundaries and physically interacts with cohesin (Parelho et al., 2008; Wendt et al., 2008)), can give rise to contact maps similar to those revealed experimentally through Hi-C (Banigan et al., 2020; Fudenberg et al., 2016; Nuebler et al., 2018). Consistent with the LEF model, recent experimental studies demonstrate that cohesin and condensin can actively drive loop formation on naked DNA (Davidson et al., 2019; Ganji et al., 2018; Haarhuis et al., 2017; Kim et al., 2019) with additional supportive evidence in vivo (Gabriele et al., 2022; Pradhan et al., 2022). Taken together, these observations suggest that cohesin and condensin are ATP-dependent drivers of dynamic DNA looping.

From a functional perspective, TADs play a critical role in defining the probability of physical contacts along the chromosome; these units correlate with chromatin attributes such as gene expression levels, suggesting that form and function of TADs are intricately linked (Ibrahim and Mundlos, 2020). It is intuitive that genetic transactions requiring interactions between DNA elements will be influenced by both TAD structure and dynamics. For example, discrete chromatin contacts that underlie critical regulatory processes such as transcriptional enhancement and co-regulation are tied to cohesin function, as is insulation and the suppression of genomic interactions with undesirable outcomes (Flavahan et al., 2016; Lupianez et al., 2015; Nora et al., 2012; Shen et al., 2012; Symmons et al., 2014). However, the static, time-averaged view of TADs arising from Hi-C models derived from thousands of cells fails to address the underlying chromatin dynamics on which TADs impinge. Moreover, in-situ hybridization approaches that can be applied to individual, fixed cells (Bintu et al., 2018; Wang et al., 2016) also cannot address how chromatin folding influences its dynamics. Indeed, how TADs, and the formation or persistence of looping interactions on which they depend, impact the dynamic properties of chromatin *in vivo* remains unclear but is a question likely to benefit from a combination of experimental and in silico approaches (Tiana and Giorgetti, 2018).

The chromosomes behave as large polymers that are crowded within the nuclear volume. Given that individual genetic loci are part of these large chromosomes and move through the crowded nucleoplasm, it is not surprising that they display a characteristic sub-diffusive behavior across model systems (Heun et al., 2001; Marshall et al., 1997; Weber et al., 2012). A distinguishing feature of the chromatin polymer is that its mobility depends on cellular energy (Heun et al., 2001; Joyner et al., 2016; Marshall et al., 1997; Weber et al., 2012). A persistent mystery is the identity of the energy-dependent process(es) underpinning this observation and whether it reflects changes in the material properties of the nucleoplasm upon energy depletion (Joyner et al., 2016) and/or an activity of an ATP-requiring enzyme(s) to agitate chromatin dynamics. Given that SMC proteins are themselves ATPases capable of translocating DNA fibers relative to each other, they are plausible candidates to be such “agitating” factors.

Here we employ a combination of live cell imaging of gene loci in fission yeast with polymer dynamics simulations to investigate how the configuration (specifically TADs) and dynamics of chromatin are linked. On short (seconds) timescales we find that several different gene loci in fission yeast display highly similar sub-diffusive dynamics regardless of location on the chromosome. Surprisingly, we find that loss of either cohesin or condensin function increases chromatin locus fluctuations, suggesting that chromatin loops made by SMC complexes predominantly constrain chromatin movement rather than driving chromatin dynamics through the act of loop extrusion. Employing Rouse-polymer simulations we independently show that it is chromatin looping that constrains chromatin mobility into a sub-diffusive state characterized by an exponent lower than the expected Rouse polymer value of 0.5, recapitulating our experimental observations. Our experimental data and simulations further argue that both the number and lifetime of loop extruding factors influence the dynamic behavior of chromatin. Last, we identify the INO80 complex as an important driver of chromatin mobility in fission yeast that likely acts in parallel with loop extrusion. Taken together, we demonstrate that energy-dependent chromatin mobility is likely driven, at least in large part, by the action of INO80, while loops, driven by cohesin and to a lesser extent condensin, introduce constraints on chromatin mobility.

## Results

### Chromatin loci show a characteristic sub-diffusive behavior in fission yeast at short timescales

To gain insights into the factors that contribute to dynamic chromatin organization *in vivo*, we developed a live-cell imaging, tracking, and analysis approach to precisely characterize the spatial and temporal dynamics of chromatin loci at short (ms-s) timescales (Fig. 1a). We visualized chromosomal loci dynamics in fission yeast by both targeted and random integrations of *lac* operator (*lacO*) repeats and expression of fluorescently-tagged *lac* repressor (GFP-LacI). Widefield fluorescence microscopy movies were obtained at short (58 ms) timesteps. The position of each individual locus at each time step was determined by fitting a Gaussian to the diffraction-limited *lacO*/GFP-LacI focus; the motion of the locus was subsequently tracked using a single particle tracking (SPT) algorithm (Crocker and Grier, 1996). Thousands of individual loci tracks were then subjected to motion analysis by calculating and fitting the mean squared displacement (MSD) versus time to theoretical profiles that take into account localization noise and motion blur (Bailey et al., 2021). Covariance analysis of the particle tracks further revealed that the motion of gene loci in fission yeast can be described by a single diffusive state (Bailey et al., 2021). On the seconds timescale, we observe a characteristic sub-diffusive behavior in fission yeast that is nearly identical at six different genomic positions (Fig. 1b and Fig. S1a). On these timescales, this characteristic chromatin motion is well described by a single mean anomalous exponent (α = 0.44 ± 0.04), which provides a good description of all experimental MSDs for the first 29 time points (between 0.058 and 1.7 s) as indicated by the best fit (solid lines) of the anomalous diffusion equation (Eq. 1, see Methods). Of note, this value is slightly but consistently lower than α = 0.5 – the value previously measured for budding yeast chromatin (Weber et al., 2012) and the expected value corresponding to the Rouse-model polymer (beads connected by springs) (Doi and Edwards, 1986). Using our approach we also recapitulate a value close to α = 0.5 for budding yeast harboring a tagged chromatin locus (Fig. S1b), suggesting that the depressed value we observe in fission yeast is likely meaningful. Plausible reasons for the difference we observe between fission and budding yeast include smaller chromosome size, differences in histone modifications (absence of histone H3 lysine 9 methylation), or TAD scale (see below). Applying an exponent of α = 0.44, the “diffusion coefficient’’, *D*, ranges from 0.0025 – 0.0031 μm^2^/s^0.44^ across the six genetic loci in fission yeast (see Supplemental Table S1). Unless otherwise noted we used α = 0.44 in all our subsequent analysis. Consistent with prior studies, depletion of ATP by the addition of sodium azide decreases the diffusivity about two-fold (from 0.0028 to 0.0015 μm^2^/s^0.44^ for the *mmf1* locus (Fig. 1c)).

**Fig. 1.**
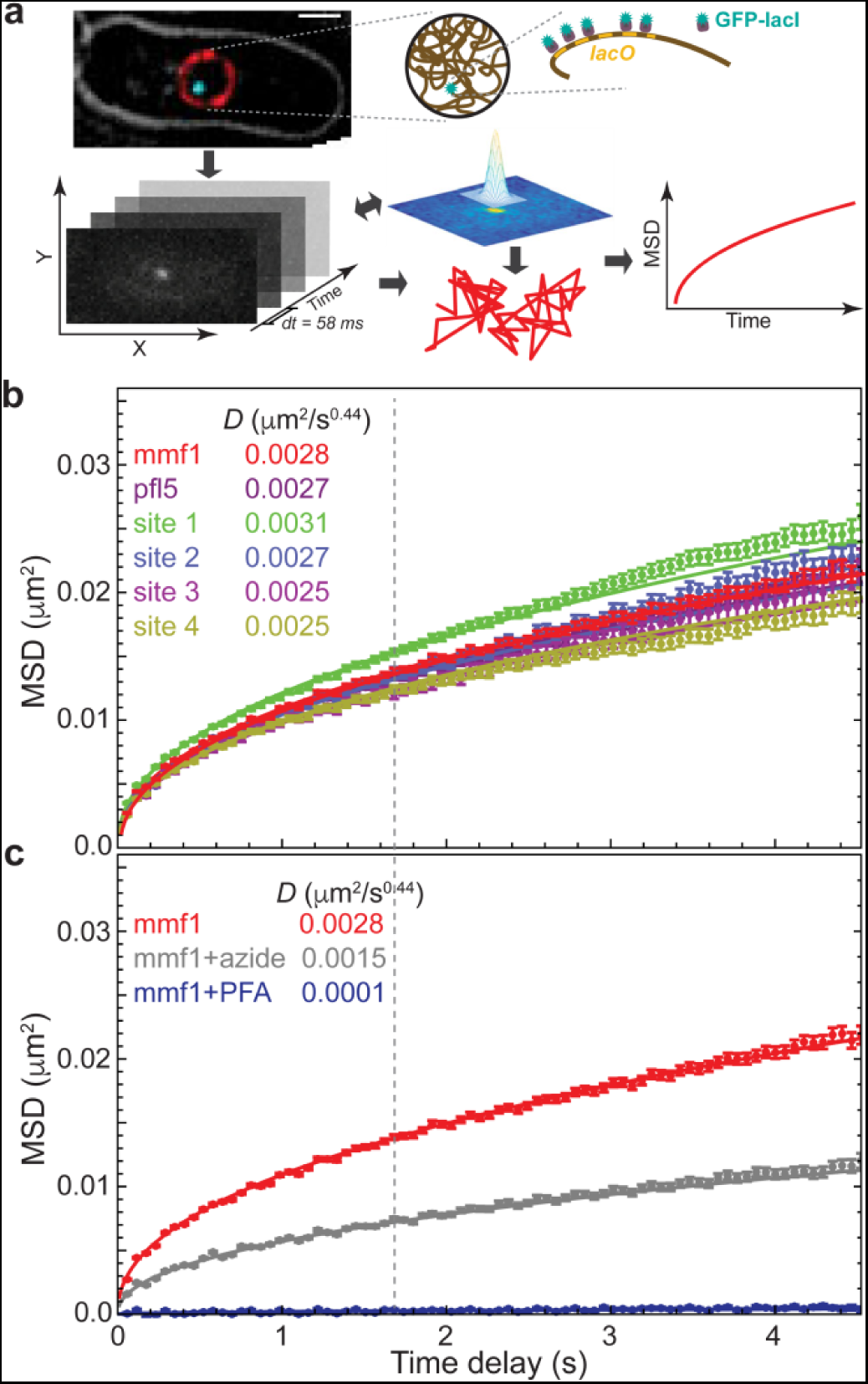
Visualization and tracking of DNA loci over time reveals characteristic, ATP-dependent chromatin dynamics in fission yeast. (**a**) Cells labeled with a lacO array inserted adjacent to the gene of interest or at random are visualized using GFP-LacI fluorescence. DNA loci are tracked using a custom 2D single particle tracking algorithm (Bailey et al., 2021). **(b)** Chromatin diffusivity is nearly identical across six different genomic locations as shown by the mean squared displacement (MSD) of each genetic locus as a function of the time. Dashed line marks “window of observation”, along with its calculated diffusion coefficient, D. **(c)** Cells depleted of ATP by treatment with sodium azide show much slower chromatin dynamics. For comparison, cells fixed with formaldehyde were imaged and analyzed to estimate systematic error in our image acquisition and analysis system. Error bars in (b) and (c) designate standard errors of the mean. Lines are the best fit of Eq.1 with indicated D values.

### Microtubule-driven centromere oscillations influence chromatin dynamics on longer time scales

In fission yeast, centromeres are mechanically coupled to the spindle pole body (SPB; the centrosome equivalent) (Funabiki et al., 1993). Thus, forces from microtubule polymerization that drive oscillation of the SPB along the long axis of the cell on the minutes timescale (Tran et al., 2001) could manifest in chromatin motion, even in regions distal to the centromere. Indeed, we observe clear super-diffusive behavior on the tens of seconds time scale (α > 1) for a *lacO* array integrated directly adjacent to the chromosome II centromere (*cen2*) (Fig. 2a). This super-diffusive motion is driven by microtubules as their depolymerization with carbendazim (methyl benzimidazol-2-ylcarbamate, MBC) leads to a strong depression of motion at *cen2* and the adoption of sub-diffusive behavior (Fig. 2a). Although more subtle, we also observe that the motion of a distant *mmf1* locus (∼ 1.8. Mb from the centromere) on the tens of seconds time scale is enhanced in cells with intact microtubule dynamics relative to cells treated with MBC (Fig. 2a, apparent after time delays of ∼5 s). We exclude that this effect is due to rigid body movement of the nucleus in response to microtubule forces exerted on the SPB on this timescale (Fig. S2). Interestingly, below the ∼5 s time delay regime *cen2* displays lower diffusivity than the *mmf1* locus (*D* = 0.0016 μm^2^/s^0.44^ in the presence of MBC), suggesting that bridging centromeres to the SPB acts to constrain chromatin motion at short times. Indeed, if we focus on the same time regime explored in Fig. 1 (up to a time delay of ∼ 4 s), microtubule dynamics play a minor role on the diffusivity of *cen2*, which is far more constrained than *mmf1* (Fig. 2b). At *mmf1* we observe only a nominal contribution of microtubule dynamics to the observed diffusivity at this time scale (*D* = 0.0026 μm^2^/s^0.44^ in the presence of MBC, Fig. 2b). We therefore chose to focus the rest of our study on this seconds-scale time regime where there is little influence of microtubule dynamics.

**Fig. 2.**
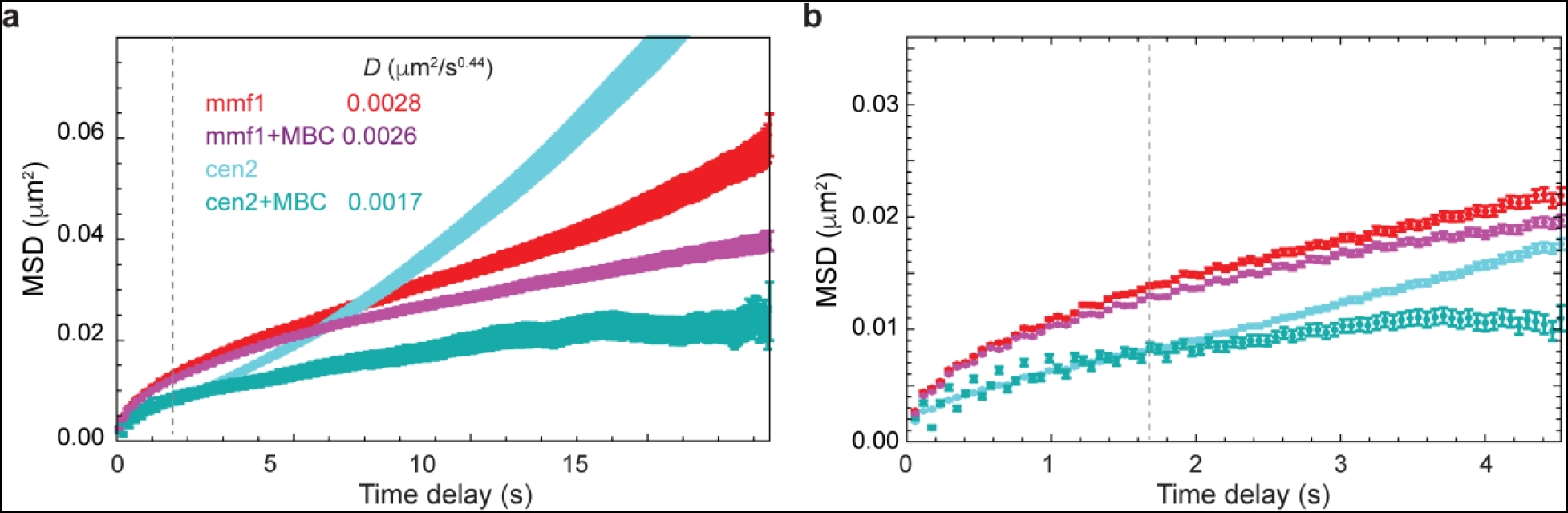
Microtubule dynamics actively drive chromatin motion at the centromeres and, to a lesser extent, the chromosome arms in fission yeast. In S. pombe, large-scale chromatin dynamics are influenced by movement of the spindle pole body through its attachment to the centromeres (Funabiki et al., 1993). (a) The centromeres (cen2) demonstrate actively-driven motion at the tens of seconds timescale as revealed by MSD analysis. Upon addition of the microtubule-depolymerizing agent carbendazim (MBC), this mobility is greatly depressed. The effect of microtubule dynamics is far less prominent for a locus in the chromosome arm (mmf1). (b) On the seconds time scale, the mobility of cen2 is far more constrained than the mmf1 locus and microtubule dynamics play a more muted effect. Error bars indicate standard errors of the mean.

### The loop extrusion complexes, cohesin and condensin, primarily constrain chromatin motion in fission yeast

Chromosome conformation is influenced by the activity of cohesin and condensin in fission yeast. Loss of cohesin strongly disrupts TADs as assessed by Hi-C (independent of its role in sister chromatid cohesion) (Mizuguchi et al., 2014) while condensin plays a more subtle but nonetheless discernible role in interphase chromatin organization (Kakui et al., 2020).

Condensin and cohesin presumably shape chromosomes through their loop extrusion activity, resulting in a dynamic steady state of DNA loops that appear and disappear in a stochastic fashion. However, prior studies in budding yeast suggest that cohesin depletion leads to an increase in chromatin diffusivity on the tens of seconds time scales (Cheblal et al., 2020; Dion et al., 2012), which was interpreted as arising due to loss of sister chromatin cohesion. As we visualize chromatin dynamics during the lengthy G2 phase in fission yeast after replication and the establishment of sister chromatid cohesion, here the critical loss of cohesin loading allows us to specifically interrogate the influence of dynamic loop extrusion on chromatin dynamics. To this end, we examined how the motion of two loci, *mmf1* and *pfl5*, which are distant from their chromosome’s centromere (∼1,800 and 2,800, kb respectively), were affected by disruption of post-replicative cohesin loading or condensin function. Specifically, we employed critical loss-of-function, temperature-sensitive alleles of the cohesin loading factor (*mis4-242*) or of the SMC2 subunit of condensin (*cut14-208*). To our surprise, we observed a profound increase in chromatin mobility (∼30 to ∼50% increase in the diffusion coefficient, *D*) upon inactivation of either loop-extruding complexes at *mmf1* (Fig. 3a), with a greater effect upon disruption of cohesin loading. These effects of disrupting LEF function on the dynamics of the *pfl5* locus were qualitatively similar (Fig. 3b). Thus, we conclude that: 1) cohesin constrains chromatin mobility independent of its role in sister chromatin cohesion, which could reflect its role in generating loops that underlies its cell cycle-independent role in TAD formation (Mizuguchi et al., 2014); and 2) condensin also contributes to constraining chromatin dynamics, albeit to a lesser extent than cohesin.

**Fig. 3.**
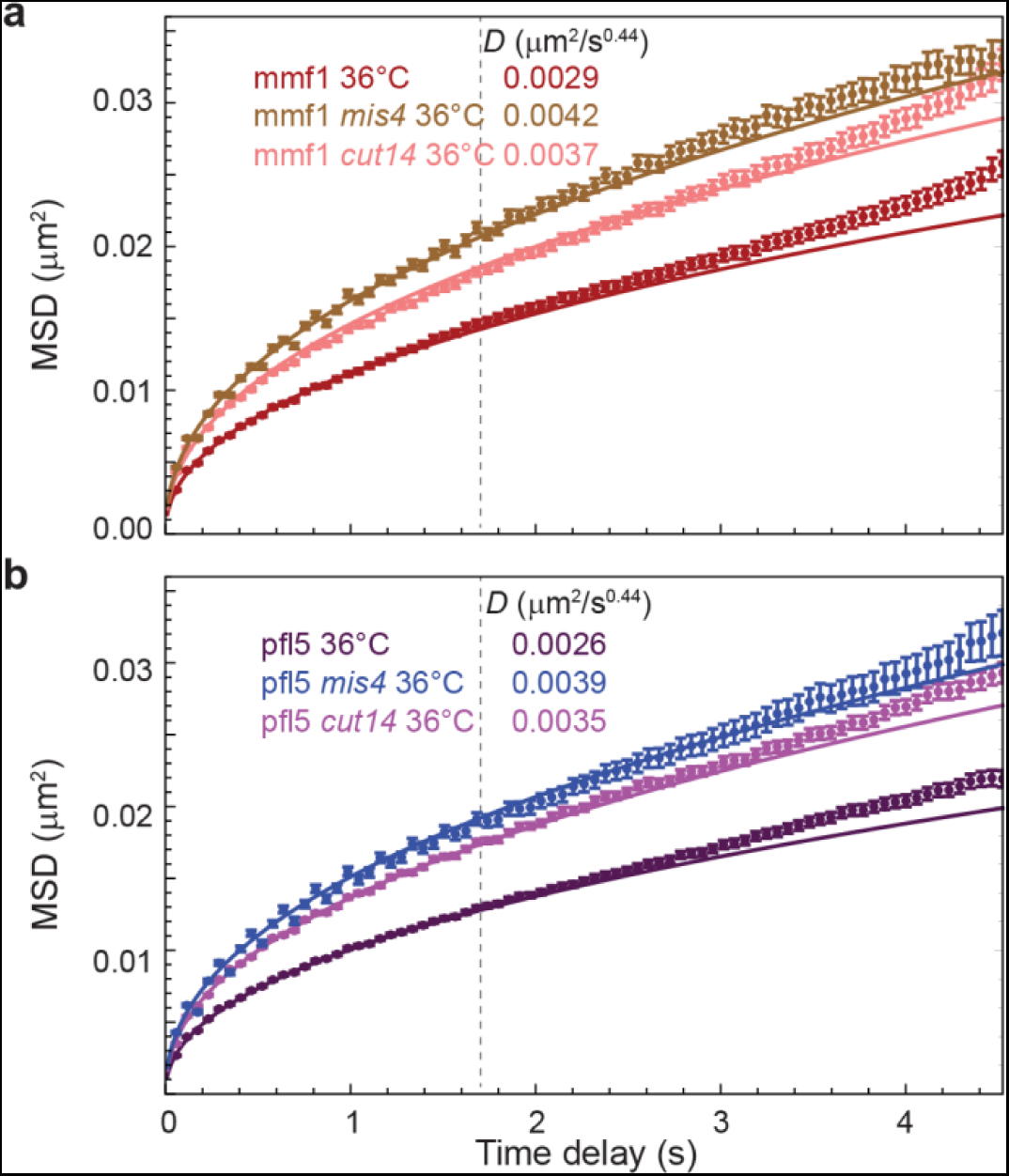
Loss of cohesin or condensin activity increases chromatin mobility. **(a)** MSD analysis of G2 cells harboring temperature-sensitive alleles of the cohesin-loading protein Mis4 and condensin complex subunit 2 (Cut14) and fluorescently labelled *mmf1* locus were imaged at the non-permissive temperature (36°C) to inhibit G2 cohesin and condensin function, respectively. **(b)** The same enhancement of chromatin dynamics is observed at a separate genomic locus, *pfl5*. Error bars indicate standard errors of the mean. Lines are the best fit of Eq.1 with indicated *D* values.

### Simulations predict that looping interactions constrain chromatin mobility

Since chromatin dynamics reflects the response of the chromatin polymer to both active processes and thermal fluctuations, we turned to modeling in order to understand how condensin- and cohesin-mediated loops arising from loop extrusion impact fluctuations of the chromatin polymer. We coupled Rouse-model polymer dynamics simulations with LEF-model simulations of the loop configuration for a given genomic region. We implemented a loop extrusion model that includes: (1) random loading of LEFs, (2) followed by bi-directional loop extrusion until a LEF encounters another LEF or a boundary element such as a site bound by CTCF (Alipour and Marko, 2012; Fudenberg et al., 2016; Nuebler et al., 2018), or (3) until the LEF dissociates, causing the loop to dissipate. As CTCF does not exist in fission yeast but CTCF and its binding sites have been reported for mouse cells, we first simulated dynamics of several 6-Mbp regions of the mouse genome. We described each 10 kb of the genome as a bead connected to its neighbors by springs of stiffness, *κ*, and experiencing a friction coefficient, *ζ*. To incorporate loops, we augmented the usual nearest-neighbor Rouse model with additional springs (of the same spring constant) that connect non-adjacent beads, thus representing the base of a loop. The loop configuration (defined by the monomers/beads that are linked) evolves according to the stochastic “LEF-CTCF” model similar to that developed by Mirny and colleagues (Fudenberg et al., 2016; Nuebler et al., 2018) as described above. In the model, CTCF occupancy determines the probability of the LEF passing through the CTCF binding sites. Using reported CTCF occupancy derived from experimental data from chromatin immunoprecipitation followed by sequencing (Bonev et al., 2017) as an input and the parameters outlined in Table S2, LEF model simulations resulted in a dynamic steady state of chromatin loops with about 30 loops within a 6 Mbp region. The comparison of MSDs between the Rouse simulations without loops (red curve) and with CTCF-dependent loops (orange curve) reveals that loops constrain chromatin motion (Fig. 4a), consistent with our experimental MSD measurements of gene loci in *S. pombe* with and without disrupted SMC complex loading or activity (Fig. 3). Increased chromatin motion was tied to chromatin conformations that were less compact in the absence of loops (example shown in Fig. 4b, red) compared to more condensed conformations in simulations with LEFs generating loops (Fig. 4b, orange).

**Fig. 4.**
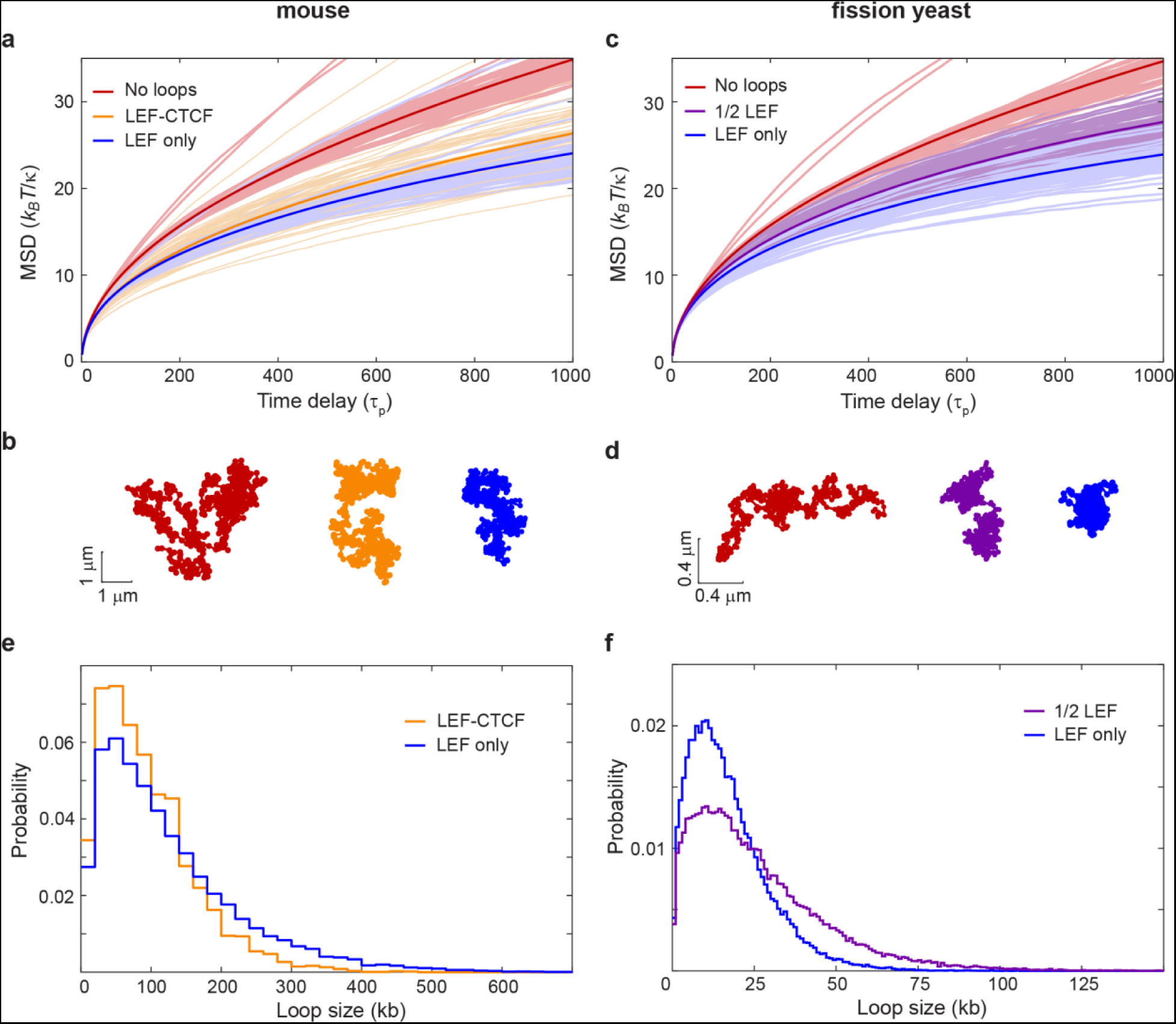
LEF activity constrains chromatin motion as revealed by Rouse-type polymer simulations. **(a)** The formation of time-dependent loops driven by loop extrusion according to the “LEF-CTCF” model with semi-permissible boundaries at CTCF binding sites (orange) or “LEF only” model lacking explicit boundaries (blue) applied to the mouse genome leads to decreased mean-squared displacements (MSD) compared to the polymer without loops (red). MSDs arise from a Rouse-model with beads of friction coefficient ζ, connected by springs of spring constant *κ* ; *τp = ζ* /(4*κ*) is the characteristic time of polymer relaxation. **(b)** Examples of the instantaneous polymer configurations from simulations for the mouse genome. **(c)** The same analysis described in (a) but applied to the fission yeast genome for the “LEF only” model (blue) compared to the polymer without loops (red). In magenta is the LEF-only model in which only half of the LEFs are present. **(d)** Examples of the instantaneous polymer configurations from fission yeast simulations. **(e)** Loop size distribution for simulations of the mouse chromatin with “LEF-CTCF” and “LEF only” models. **(f)** Loop size distribution for simulations of the fission yeast chromatin with “LEF only” models at “full” or “1/2” numbers of LEFs.

We next developed an analogous simulation approach that could be applied to the fission yeast genome. Yeasts lack CTCF, although they likely have alternative boundary elements that define experimentally observed TADs (Mizuguchi et al., 2014; Thon et al., 2002). We therefore devised a simulation, which we call the “LEF only” model, that includes: 1) random loading of LEFs and 2) the ability of the LEF to move bi-directionally from its loading site until it encounters another LEF, (3) or until it dissociates – a feature of many prior loop extrusion models (Alipour and Marko, 2012; Fudenberg et al., 2016; Nuebler et al., 2018). The “LEF only” model is the same as the “LEF-CTCF” model with one exception – there is no influence of CTCF (or any other explicit factor) on LEF movement. Similar to the CTCF-dependent mouse genome results, we again observe that LEF activity leads to a reduction in the MSD in simulations of a 300-kb region of the fission yeast genome (Fig. 4c, compare the blue curve to the red) and more compact polymer conformations (Fig. 4d in blue, compare to without loops, in red). Moreover, subjecting the mouse genome to this alternate LEF-only loop extrusion model also constrains loci motion (Fig. 4a-b, blue curve and polymer conformation). The resulting MSDs in the LEF- only model are suppressed slightly beyond those generated by the LEF-CTCF model (Fig. 4a), which is also reflected in more compact polymer conformations (Fig. 4b). Higher compaction stems from the lack of CTCF-dependent pausing sites, which allows LEFs to travel more genomic distance before encountering another LEF, leading to larger chromatin loops on average (Fig. 4e), and therefore a smaller fraction of chromatin existing outside of loops. These results suggest that the distribution of loop sizes can impact chromatin dynamics.

We next investigated how the number of loops impacts the MSD. If we decreased the number of LEFs by half (referred as “1/2 LEF” model) in our fission yeast simulation we observed an MSD profile and polymer compaction that is intermediate between the no-loop conformations and the “full amount” of LEFs (Fig. 4c, purple curve). This simulation result can, at least in part, explain the observed effect that inactivating cohesin or condensin has on loci motion (Fig.3), as loss of either increases chromatin mobility despite the continued presence of the other presumed LEF. Indeed, while having fewer SMCs allowed formation of larger loops (reflecting a reduced probability of encountering another SMC during the LEF residence time), the extent of polymer compaction in the “1/2 LEF” model was also intermediate, as a greater fraction of the chromatin was outside of loops (Figs. 4d, 4f). Taken together, these observations again reinforce the trend that loop size and density are important variables of how chromatin conformation impacts its dynamics. Moreover, our observations underscore that while particular LEF models might be different in their details (i.e. with or without CTCF), extruded loops consistently constrain chromatin mobility.

### The long residence time of cohesin likely underlies its dominance over condensin in constraining chromatin mobility

Our experimental data suggest that cohesin has a more profound influence on constraining chromatin dynamics than condensin given that disrupting its loading in G2 cells quantitatively drives up the MSD to a greater extent than loss of condensin function (Fig. 3a-b). While our 1/2 LEF simulation provides an explanation for the sensitivity of chromatin mobility to loss of either cohesin or condensin, it does not explain why they have different impacts on loci dynamics. We hypothesized that the differential effect could stem from distinct lifetimes of cohesin and condensin on chromatin, as supported by prior experimental observations (Gerlich et al., 2006a; Gerlich et al., 2006b; Hirano, 2016). To begin to explore this hypothesis, we carried out simulations in which we explicitly considered two types of LEFs differing 10-fold in their respective lifetimes while keeping the total number of LEFs constant. As expected, introducing activities of both long- and short-lifetime LEFs (mimicking cohesin and condensin, respectively) resulted in constrained MSDs (Fig. 5a and Fig. S3a, blue versus red curves), similar to those in previous simulations (Fig. 4c). Here, however, inactivation of long-lived LEFs (Fig. 5a and Fig. S3b; dark green) led to a much stronger mobility enhancement compared to the inactivation of short-lived LEFs (Fig. 5a and Fig. S3a; light green). Thus, LEFs with shorter lifetimes have a weaker impact on chromatin dynamics, perhaps because the loops they form during their lifetime are smaller (Fig. S3c). By contrast, LEFs with longer lifetimes make larger loops (Fig. S3c) that may explain why they dominate in compacting chromatin and, as a consequence, constraining chromatin mobility.

**Fig. 5.**
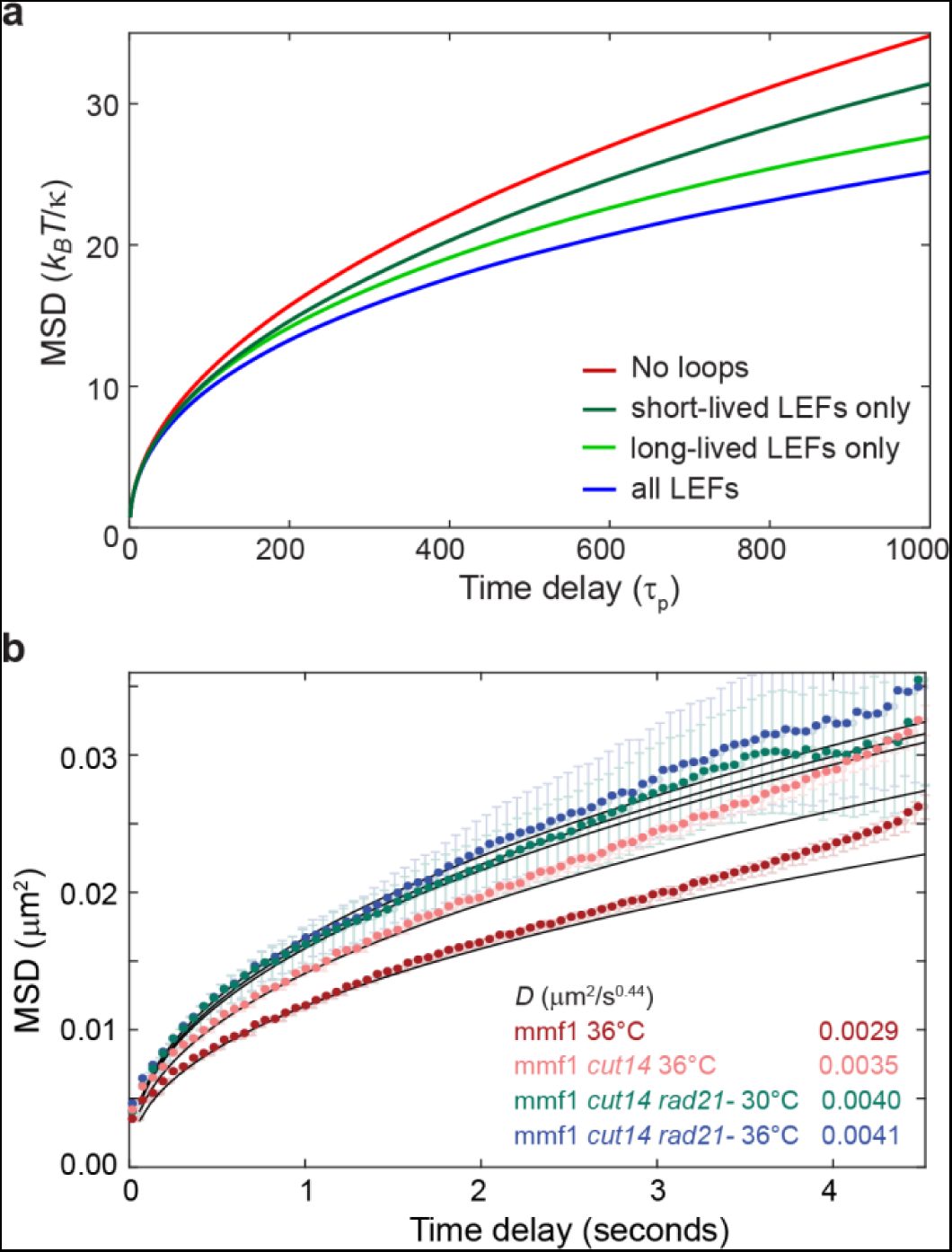
Impact of loop extrusion on chromatin mobility depends on the LEF lifetime. **(a)** MSD results of the Rouse-type polymer simulations combined with loop-extrusion simulations that considered two types of LEF complexes (in equal amounts) with different lifetimes. Blue – simulations with both types of LEFs present, green – only long-lived LEF present, dark green – only short-lived LEF present, red – no LEFs. Lines represent the average over all beads and all simulations. See also Fig. S3. **(b)** Experimental MSDs for WT cells at 36°C (red), cells harboring a temperature-sensitive allele of the condensin complex subunit 2 (Cut14) imaged at the non-permissive temperature (36°C, pink), cells harboring the temperature-sensitive allele of Cut14 imaged after depletion of the Rad21 subunit of the cohesin complex at the permissive temperature (30°C, green) and at the non-permissive temperature (36°C, blue). Lines are the best fit of Eq.1 with indicated *D* values.

To further examine the relative contributions of cohesin and condensin to chromatin mobility experimentally we engineered a strain that combines critical Rad21 depletion (a structural component of cohesin) with the temperature-sensitive *cut14-*208 allele in condensin. While rapid (∼20 min, Fig. S4) Rad21 degradation at the permissive temperature for *cut14-*208 led to enhanced *mmf1* locus mobility (Fig. 5b) similar to inactivation of Mis4 (Fig. 3a), additional inactivation of Cut14 by shifting to its non-permissive temperature had only a nominal effect.

These results suggest that cohesin is the dominant factor responsible for damping chromatin dynamics, plausibly due to its longer lifetime. Taking our experimental and simulation findings together suggests that, on this time scale, the loop-extruding activity of cohesin, and to a lesser extent condensin, constrains the MSD of chromatin, which is sensitive to both the number and size of loops.

### Loop extrusion decreases the effective exponent describing chromatin diffusivity

We next inquired whether the loop configuration simulations could provide further insight into our observation that the sub-diffusive chromatin motion in fission yeast is described by an exponent, α, of ∼ 0.44, which deviates from that expected for a Rouse polymer (α = 0.5). To this end, we calculated the effective exponent, α = [log MSD(t_n+1_)−log MSD(t_n_)]/[log t_n+1_−log t_n_] from the simulated chromatin configurations and compared how α is influenced by loop extrusion in the simulations. As expected, the effective exponents for the classical Rouse model lacking loops in mouse simulations lies close to 0.5 for the wide range of times longer than the polymer relaxation time *τ_p_*, *τ_p_ = ζ*/(4*κ*) (Fig. 6a, red curve). Remarkably, however, the effective exponent in the presence of loop extrusion employing either the CTCF-LEF model (Fig. 6a, orange curve) or the “LEF-only” model (Fig. 6a, blue curve) shows a gently evolving value with a mean that appears to vary continuously from a value near α ≃ 1/2 for *t* ≃ *τ_p_* to a value near α = 0.4 for *t* ≃ 10^3^*τ_p_*. Similarly, loop extrusion depresses the value of the effective exponent in the fission yeast “LEF only” or “1/2 LEF” simulations (Fig. 6b). These results are strikingly similar to our experimental finding that MSDs in *S. pombe* are well-described overall by a value of α ≃ 0.44 (Fig. 1, Table S1). Thus, our data support a model in which chromatin looping interactions not only influence its apparent diffusivity, but also manifest in depressed values for α compared to that of a Rouse polymer.

**Fig. 6.**
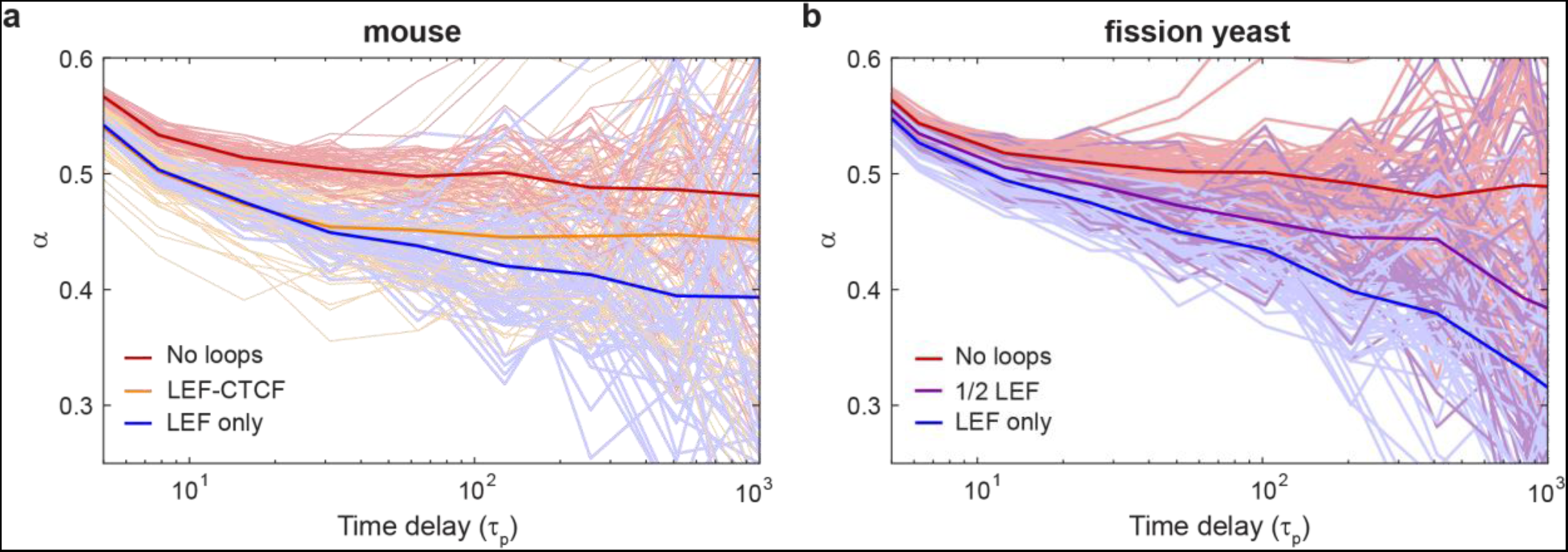
Loop extrusion decreases the apparent sub-diffusive exponent. **(a)** and **(b)** Effective “instant” exponent, α, versus time for the mouse and fission yeast models presented in Fig. 4. Thin lines correspond to individual beads, thick lines represent the average over all beads and simulations.

### The INO80 chromatin remodeler drives chromatin mobility in fission yeast

Both our experimental and simulation approaches find that loop extrusion by SMC complexes predominantly suppresses chromatin mobility rather than serving as an activity that promotes its dynamics. Nonetheless, since ATP-depletion drastically reduces chromatin fluctuations and motion (Fig. 1c), there must be a process(es) tied to cellular metabolism that act on the chromatin polymer to elicit its fluctuations. ATP-dependent chromatin (nucleosome) remodelers represent attractive alternative candidates for driving energy-dependent chromatin mobility and have been previously implicated in driving enhanced chromatin mobility in response to DNA damage (Cheblal et al., 2020; Hauer et al., 2017; Neumann et al., 2012; Seeber et al., 2013).

Whether this activity is also tied to LEF function, and more generally if and how nucleosomes impact on the translocation of LEFs along the DNA, remains incompletely understood. To test whether chromatin remodelers contribute to ATP-dependent chromatin mobility in fission yeast, we examined cells lacking Arp8, an auxiliary component of the essential INO80 remodeling complex, and Arp9, a component of both the SWR1 and RSC remodeling complexes. Strikingly, loss of Arp8 led to a clear suppression of chromatin mobility (*D* = 0.0024 μm^2^/s^0.44^), representing a 40% decrease in the ATP-dependent diffusivity, i.e., above the basal, ATP-depleted motion (Fig. 7a). By contrast, loss of Arp9 had little effect (Fig. 7a). Importantly, as INO80 is essential for viability, Arp8 deletion is akin to a hypomorphic allele rather than a complete loss-of-function, leaving open the possibility that INO80 activity contributes to ATP-dependent chromatin diffusivity to an even greater extent than is observed in *arp8Δ* cells. These results are consistent with a prior study in which INO80 (but not SWR1 or RSC) was both required for a transcription-dependent boost in chromatin mobility and was sufficient to impart increased diffusivity when artificially targeted to a chromosomal locus in budding yeast (Neumann et al., 2012), although the unique role for INO80 but not SWR and RSC remains unexplained.

**Fig. 7.**
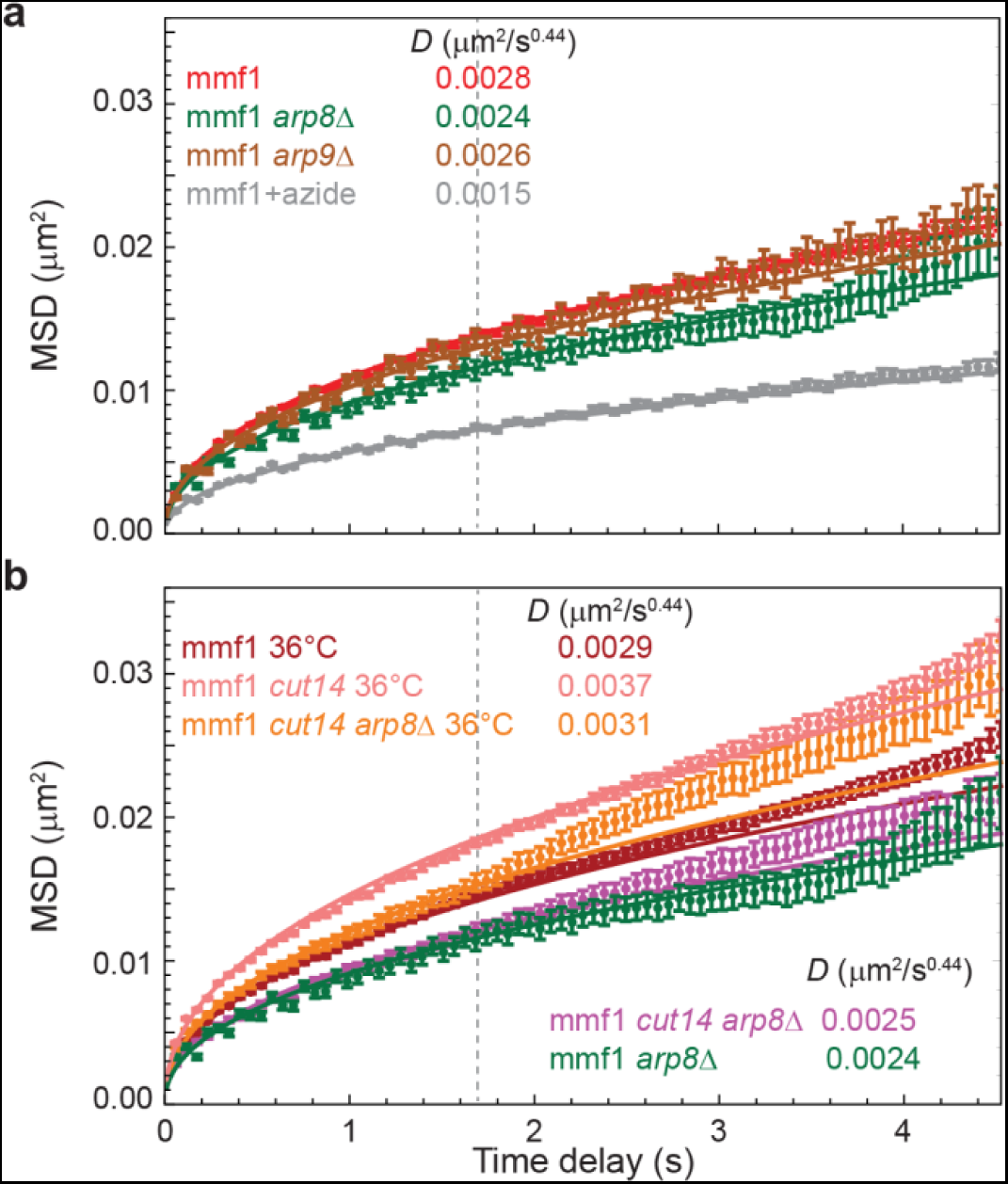
Loss of the nucleosomal remodeling complex protein Arp8 decreases chromatin mobility likely through a mechanism distinct from loop extrusion. (a) Loss of Arp8, a component of the INO80 complex, reduces ATP-dependent chromatin mobility by ∼40%, while loss of Arp9, a component of both the SWR1 and RSC complexes, has minimal effect. For comparison, cells depleted of ATP (treated with sodium azide) are shown (replotted from Fig. 1c). (b) Loss of Arp8 dampens the elevated chromatin dynamics in the cut14-208 background at the non-permissive temperature to a slightly less extent than in WT cells (a ∼30% decrease) but remains much more diffusive than at the permissive temperature. Error bars indicate standard errors of the mean. Lines are the best fit of Eq.1 with indicated D values.

To address the interplay between the boosting of chromatin mobility by INO80 and the constraint imposed by LEFs, we examined how loss of Arp8 and the elevated chromatin diffusivity observed in LEF mutants would intersect. At the permissive temperature, *cut14-208 arp8Δ* cells displayed decreased diffusivity of the *mmf1* locus similar to *arp8Δ* cells at the short times used in our analysis, although there is a slight upwards trend at longer times in cells also harboring the *cut14-208* allele (Fig. 7b, *D* = 0.0025). At the non-permissive temperature (36°C), loss of Arp8 also dampens the elevated chromatin dynamics observed in the *cut14-208* background (Fig. 7b; *D* = 0.0031 μm^2^/s^0.44^). The relative magnitude of the effect (a ∼30% decrease of the ATP-dependent diffusivity; Fig. 7b) is only slightly smaller than that observed at the permissive temperature or upon Arp8 deletion in WT cells. Importantly, the level of absolute diffusivity in *cut14-208 arp8Δ* cells at the non-permissive temperature remains substantially elevated compared to cells lacking only Arp8 (*D* = 0.0031 versus 0.0024 μm^2^/s^0.44^). Taken together, these observations suggest that 1) INO80 activity is a major driver of non-thermal chromatin fluctuations in fission yeast and 2) loop extrusion influences the manner in which these fluctuations act on the chromatin polymer.

## Discussion

Here we provide new insights into the relationship between chromosome structure and dynamics. Combining live cell imaging and polymer simulations we converge on the conclusion that chromatin loops, which arise by SMC-driven loop extrusion, primarily constrain chromatin mobility. This constraint manifests as a decrease in the MSD of chromatin loci and the effective exponent describing their sub-diffusive behavior. We provide evidence that the activity of the INO80 chromatin remodeler is a major source of the as-yet mysterious, energy-dependent activity that drives chromatin motion. Our findings emphasize that both the chromatin conformation and the molecular machines acting on the polymer together determine chromatin dynamics.

### Loops as constraints to chromatin mobility

We observe that cohesin and condensin, both harboring ATPase activity, constrain chromatin mobility *in vivo*; a similar observation was recently reported for the effect of cohesin depletion in mouse ES cells (Mach et al., 2022). These data further mirror prior observations in budding yeast for cohesin depletion (Cheblal et al., 2020), although this effect was ascribed primarily to an effect on sister chromatid cohesion rather than loop extrusion. Our polymer simulations provide a mechanistic explanation for the constraining effect of loop extrusion on chromatin dynamics. Specifically, while loop extrusion can drive changes in the loop configuration, the “polymer” (chromatin) relaxation time scale through which this change manifests is much shorter (on the sub-second time scale) than our observations of chromatin dynamics (on the seconds timescale). Thus, changes to an explicit loop configuration have less impact on chromatin dynamics than the statistical impact that loops generally exert on the experimentally observed MSD and exponent of α = 0.44, which when related to the simulations in Fig. 6b provides a window into the values of τp relevant to our measured, second-time scale trajectories. This interpretation is further bolstered by the observation that motion of individual loci is mostly independent of their genomic position (Fig. 1b). Taking a different simulation approach for examining the effect of loop extrusion that also accounts for volume exclusion, Mach et al. likewise found recently that it is the act of loop extrusion rather than the influence of specific boundaries (or barriers) that impacts polymer dynamics, although they did not observe changes to the anomalous exponent, α (Mach et al., 2022). Taken together, these observations highlight that chromatin dynamics is dominated by the polymer nature of the chromatin and not by the local genomic conformation on the seconds time scale.

One important ramification of this model is that it provides another, “dynamic” means by which loop extrusion factors can antagonize inter-TAD interactions and reinforce the relative over-representation of local, intra-TAD interactions that extends beyond roles in imposing explicit TAD boundaries. This effect would also be expected to disfavor longer-range, stochastic interactions, consistent with Hi-C studies demonstrating that cohesin antagonizes compartments (Gassler et al., 2017; Rao et al., 2017), suggesting that loss of loops leads to more unfettered contacts between self-associating chromatin landmarks, for example those driving B compartment cohesion (Falk et al., 2019). We expect that the influence of loops on chromatin dynamics is likely to play a fundamental and conserved role as it is independent of the specific mechanisms determining the position of TADs across eukaryotic models (e.g. organisms outside of bilateria that lack CTCF and/or employ other insulator proteins such as the widely conserved TFIIIC complex (Van Bortle and Corces, 2012)). Indeed, the effect of loops on chromatin dynamics could easily extend beyond cohesin and condensin, for example to so-called tethering elements that mediate non-SMC loops (Batut et al., 2022).

### The ATP-dependence of chromatin mobility and nucleosome remodelers

The source of ATP-dependent chromatin mobility has been a perennial mystery, ever since the first live-cell recordings of chromatin dynamics were made using the *lacO-*LacI technology over twenty years ago (Heun et al., 2001; Marshall et al., 1997). Importantly, a study in budding yeast (and extended to fission yeast) demonstrates that caution must be exercised when interpreting experiments that disrupt cellular energy by glucose starvation, which results in altered cell volume, as this can lead to increased crowding that affects not just chromatin mobility but also other large macromolecular complexes (Joyner et al., 2016); for this reason, we have used sodium azide treatment to uncouple effects of ATP depletion from altered crowding in this study.

A renewed interest in actively-driven chromatin mobility has arisen through studies of the DNA damage response across multiple model systems (Zimmer and Fabre, 2019). Induction of a DNA double-strand break (DSB) leads to an increase in mobility of not only the broken chromosome region, but, surprisingly, the entire genome (Lawrimore et al., 2020; Miné-Hattab and Rothstein, 2013); this effect has been suggested to promote the homology search phase of DSB repair. While many factors contribute to this response, including nuclear and cytoplasmic cytoskeletal proteins (Caridi et al., 2018; Lawrimore et al., 2020; Lottersberger et al., 2015; Oshidari et al., 2018; Schrank et al., 2018; Swartz et al., 2014; Zhurinsky et al., 2019), of note the INO80 nucleosome remodeler appears to play a central role (Cheblal et al., 2020; Hauer et al., 2017; Neumann et al., 2012; Seeber et al., 2013). Indeed, in budding yeast loss of Arp8 disrupts the observed DSB-dependent boost in chromatin mobility (Cheblal et al., 2020; Hauer et al., 2017; Neumann et al., 2012; Seeber et al., 2013). Although the potential involvement of INO80 in multiple steps of DSB repair complicates the interpretation of the mechanisms at play, it nonetheless underscores the specificity underlying the unique relationship between the INO80 nucleosome remodeler and chromatin dynamics, possibly due to INO80’s role in nucleosome eviction (Cheblal et al., 2020).

Lastly, transcription has also been observed to drive a boost to chromatin dynamics across model systems (Chuang et al., 2006; Gu et al., 2018; Neumann et al., 2012). Importantly, in budding yeast, the increase of chromatin mobility upon transcriptional activation again requires INO80 activity (Neumann et al., 2012). Further, in perhaps the most elegant demonstration of sufficiency, simply locally targeting INO80 to a *lacO* array is sufficient to recapitulate the transcription-driven boost in chromatin mobility even in the absence of transcription; this is not the case for other nucleosome remodeling complexes when similarly targeted (Neumann et al., 2012). Thus, a common theme is that a unique function of INO80 among nucleosome remodeling complexes lies at the heart of enhanced chromatin motion in response to DSBs or transcriptional activation. Our observations suggest that a critical role for INO80 in promoting chromatin mobility extends beyond such specialized contexts and is instead generalizable, driving a component of the ATP-dependent motion characteristic of chromatin.

### Crosstalk between loop extrusion, nucleosomes, and chromatin remodelers

The potential role that nucleosome remodelers play in facilitating loop extrusion remains to be fully investigated, as we are just beginning to define mechanistically how nucleosomes impact loading and translocation of cohesin and condensin, particularly in living mammalian cells.

Single-molecule experiments show that nucleosomes impede cohesin translocation along DNA (Stigler et al., 2016) and that nucleosome removal promotes efficient loop extrusion in *Xenopus* egg extracts (Golfier et al., 2020). Moreover, numerous studies suggest that cohesin loading on chromosomes occurs at nucleosome-depleted regions (often associated with transcription start sites) and requires ATP-dependent nucleosome remodeling activities (Garcia-Luis et al., 2019; Golfier et al., 2020; Munoz et al., 2019). In addition, it has been suggested that cohesin translocation requires transcription-coupled nucleosome remodeling (D’Ambrosio et al., 2008; Dubey and Gartenberg, 2007; Glynn et al., 2004; Hu et al., 2011; Lengronne et al., 2004; Ocampo-Hafalla et al., 2016; Schmidt et al., 2009). However, recent studies suggest the possibility that loop extrusion can proceed over nucleosomes, albeit at low nucleosome density (Golfier et al., 2020; Kim et al., 2019) and/or that loop extrusion may not require topological engagement at all, allowing blockades such as nucleosomes to be circumvented (Pradhan et al., 2022). If nucleosomes (or a subset of nucleosomes, if histone modifications have an impact) do serve as barriers to loop extrusion, then ATP-dependent nucleosome remodeling would be expected to play important roles in TAD formation. Our work suggests that the INO80 chromatin remodeler carries out an important role in driving chromatin mobility in a manner that appears independent of the impact of looping interactions on chromatin dynamics. Of note, critical depletion of INO80 in budding yeast had a less pronounced effect on contact probability maps than depletion of other nucleosome remodelers (Jo et al., 2021), consistent with a loop extrusion-independent function. However, this does not rule out collaboration of nucleosome remodelers on the act of loop extrusion, which remains an exciting avenue for further study.

---

**Fig. S1.**
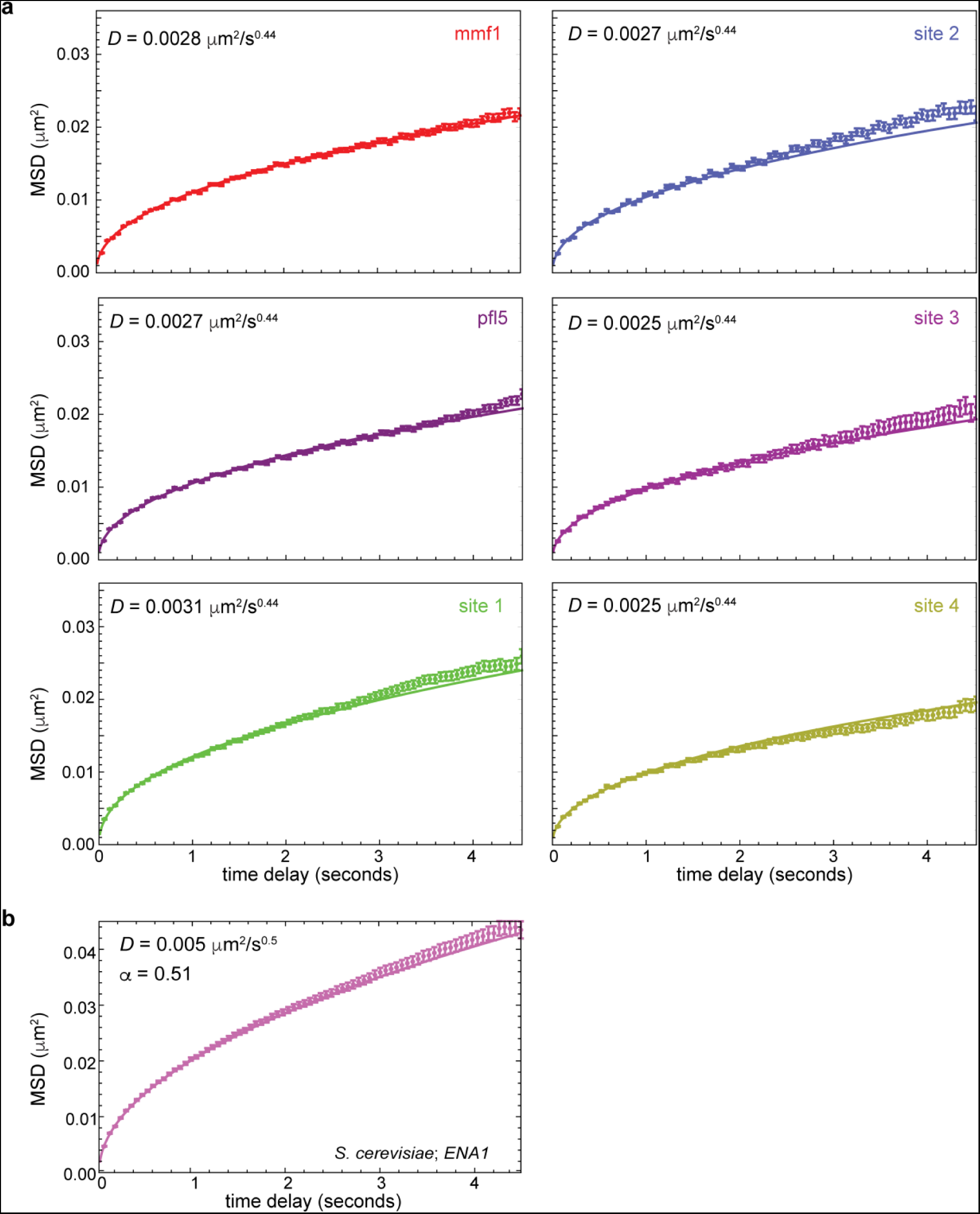
*Individual MSD plots for fission yeast and budding yeast.* (a) Chromatin diffusivity is nearly identical across six different genomic locations as shown by the mean squared displacement (MSD) of each genetic locus as a function of the time window of observation, along with its calculated diffusion coefficient, *D*. **(b)** For comparison to previous chromatin dynamics measurements, we performed our visualization/tracking/diffusive analysis regime on *S. cerevisiae* cells integrated with a *lacO* array at the *ENA1* locus, resulting in a comparable diffusion coefficient as reported previously. Error bars indicate standard errors of the mean.

**Fig. S2.**
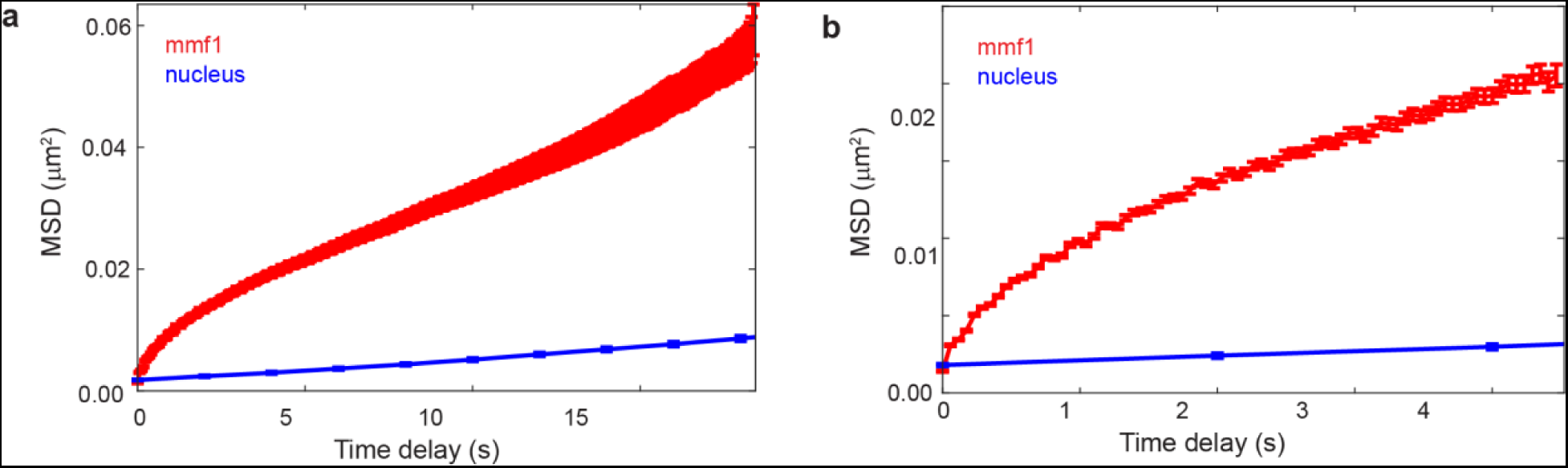
Microtubule dynamics-driven chromatin motion cannot be explained by whole-nucleus motion. **(a)** and **(b)** MSD of *lacO* array near *mmf1* relative to MSD of the whole-nucleus motion at different timescales.

**Fig. S3.**
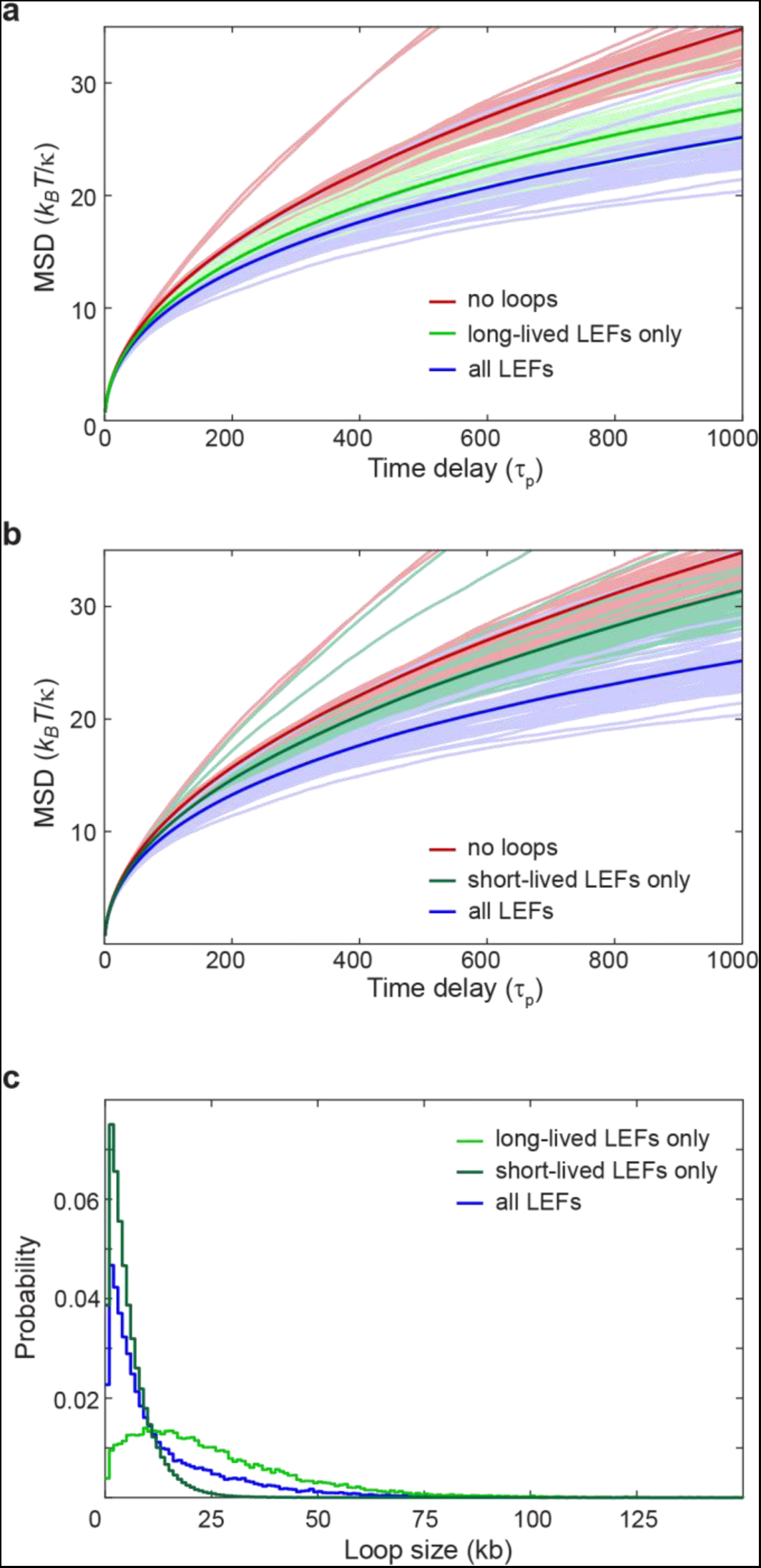
Impact of LEFs on chromatin mobility depends on LEF lifetime(s). As in Fig. 5a but showing MSDs for individual beads split into two panels for clarity. **(a)** MSD results of the Rouse-type polymer simulations combined with loop-extrusion simulations that considered two types of LEF complexes (in equal amounts) with different lifetimes. Blue – simulations with both types of LEFs present, green – only the long-lived LEFs, red – no LEFs. Thin lines represent MSDs for each individual bead, thick lines represent the average over all beads and all simulations. **(b)** As for (a) but for simulations with only short-lived LEFs (dark green) compared to simulations with both LEFs (blue) and no LEFs (red). **(c)** Loop size distribution for simulations of fission yeast chromatin subjected to the “2 LEFs” model with both LEFs present (blue), only the long-lived LEFs present (green), and only the short-lived LEFs present (dark green).

**Fig. S4.**
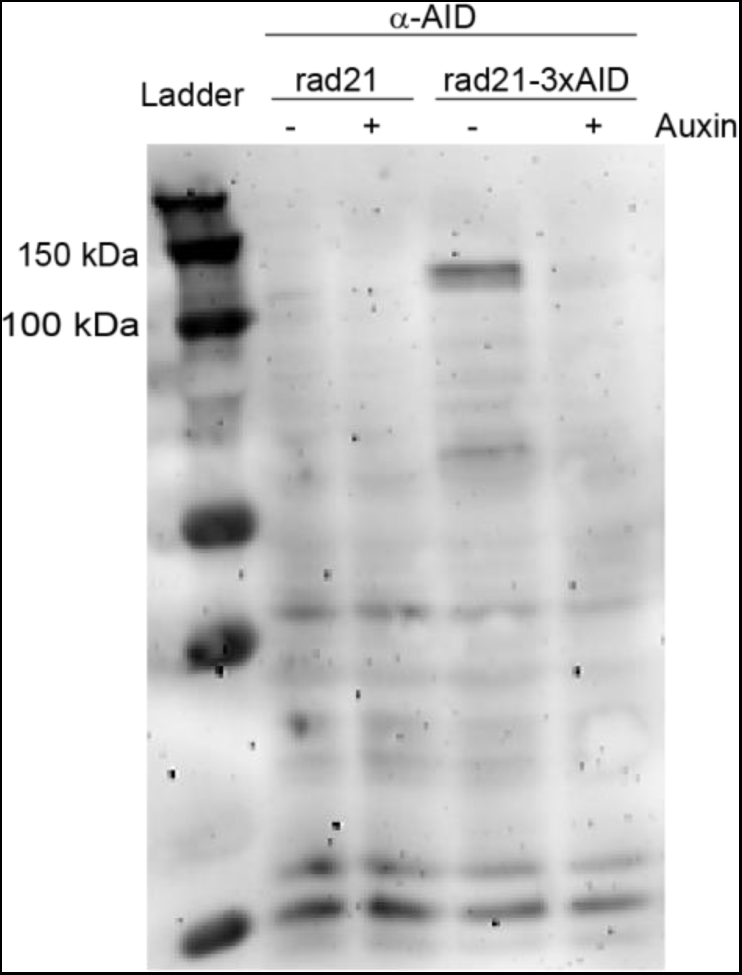
Rad21-AID can be quickly depleted from cells upon addition of the auxin analogue, 5-IAA. Western blots of lysates of WT cells or cells with Rad21 tagged with auxin-inducible degron before or after 5-IAA treatment for 20 min.

## Materials and Methods

### Strain generation and culturing

All strains used in this study are listed in Table S3. *S. pombe* were grown, maintained, and crossed using standard procedures and media (Moreno et al., 1991). Gene replacements were made by exchanging open reading frames with various MX6-based drug resistance genes (Bähler et al., 1998; Hentges et al., 2005). Targeted *lacO* array insertions were generated using a modified two-step integration procedure that first creates a site-specific DNA double-strand break to increase targeting efficiency of the linearized plasmid pSR10_ura4_10.3kb (Leland and King, 2014); integration sites are listed in Table S3. Random *lacO* array insertions were generated by transformation of the GFP-LacI-expressing strain MKSP1120 with linearized plasmid pSR10_ura4_10.3kb followed by selection on plates lacking uracil as in (Leland and King, 2014). Strains that had successfully inserted *lacO* array were identified by fluorescence microscopy. Strains with auxin-inducible degron (AID) system were constructed as follows. MKSP3626 and MKSP3629 were generated by C-terminally tagging *rad21* with *XTEN17-3xsAID-KanMX* cassette in DY48569 (Zhang et al., 2022) and in MKSP2760, respectively. DY48569 was provided by the Yeast Genetic Resource Center (NBRP, Japan). The *XTEN17-3xsAID-KanMX* cassette was obtained by amplifying from pDB4581 plasmid (a gift from Li-Lin Du, Addgene, plasmid #171124) with the megaprimers that were generated using isolated wild type *S. pombe* genomic DNA as a template and primer pairs

KL101 (GATTCACTTTTTGACGCTCCTCC) - KL102 (gttaattaacccggggatccgTAGTGATGAAAGTAGCATTCCACGTTTA) and

KL103 (agtttaaacgagctcgaattcatcgGAGGTCGGTTAATATTTTTTCAAAATCCAATTAGATCTAT)

- KL104 (GATCAATCATTGAGAATAAATTAAAAAGCGCGT). Strain MKSP3652 and MKSP3660 are two clones created by crossing MKSP3629 with DY48569. The *S. cerevisiae* strains used in this study were constructed, grown and imaged as described (Colombi et al., 2018).

### Microscopy

Cells were grown in YE5S media (yeast extract plus five supplements; (5 g/L yeast extract (BD Bacto), and 0.25 g/L adenine, 0.25 g/L histidine, 0.25 g/L leucine, 0.25 g/L lysine, and 0.25 g/L uracil, (Sunrise Science Products)) at 30°C to log phase (OD600 0.5–0.8). To fix cells, cells were incubated with 2% paraformaldehyde for 10 min and washed once with 2X PBS (phosphate-buffered saline). For temperature-sensitive alleles, the cells were grown at 30°C then shifted for twenty minutes to the non-permissive temperature (36°C) prior to imaging. For pharmacological inhibition studies, cells were incubated at 30°C in YE5S supplemented with sodium azide (0.02% w/v) or with 50 μg/ml MBC (methyl-2-benzimidazol-2-ylcarbamate) for 10 min prior to imaging and then imaged on agarose pads (see below) containing the drug at the same concentration. For auxin-induced degradation, cells were imaged on agarose pads (see below) containing 100nM 2-[5-(Adamantan-1-yl)-1*H*-indol-3-yl]acetic acid (“5-IAA”; TCI Product number A3390).

For imaging, 1.4 μL of the concentrated cell suspension was transferred to a ∼1 cm x 1 cm x 1 mm pad made of 1.4% agarose (Denville Scientific) in EMM5S media (Edinburgh minimal media plus five supplements 0.25 g/L adenine, 0.25 g/L histidine, 0.25 g/L leucine, 0.25 g/L lysine, and 0.25 g/L uracil (Sunrise Science Products)) on a microscope slide. The cell pad was covered with a #1.5 – 22 x 22 mm coverslip, and edges were sealed with VALAP (1:1:1 petroleum jelly, lanolin, and paraffin) to limit evaporation during imaging.

Fluorescence and bright field images were acquired on a DeltaVision widefield microscope (Applied Precision/GE) equipped with a temperature control chamber, 1.4 NA 100x objective (Olympus), solid-state-based illumination (Lumencor), and an Evolve 512 EMCCD camera (Photometrics). Fluorescence was excited at 488 nm and collected with emission filters between 500-550nm. Typically, 1000 images were continuously acquired with 10 ms exposure time and 58 ms per frame rate.

### Western blot

To measure rad21-XTEN17-3sAID degradation, cells were grown exactly as for imaging experiments, in YE5S media to OD = 0.6, and split in two samples – with and without addition of 100nM 5-IAA for 20 min. Cultures were centrifuged at 2000× *g* for 5 min and cell pellets were washed with 1 mM EDTA. Cells were then pelleted and lysed in 2M NaOH incubated for 10 min on ice. An equal volume of 50% trichloroacetic acid (TCA) was mixed in before incubating for 10 min on ice and collecting the protein precipitate by centrifugation. The pellet was then washed with -20°C acetone and air-dried for 15 min. The pellet was then dissolved in 5% SDS followed by an equal volume of SDS-PAGE sample buffer containing urea (24 mM Tris-Cl pH 6.8, 9 M urea, 1 mM EDTA, 1% SDS, 10% glycerol). Samples were then shaken at 37°C for 15 min before being centrifuged at 14,000× *g* for 15 min. Approximately equal loads of extracted protein were subjected to SDS-PAGE and transferred to a 0.2 µm nitrocellulose membrane (Bio-Rad 1620112). Transferred proteins were then stained with Ponceau S Solution (Sigma-Aldrich P3504). Blots were blocked (5% (w/v) dry milk/TBST) prior to incubation with primary mouse α-mAID antibody (MBL M214-3) (diluted 1:1000 in blocking buffer) overnight. The blots were then incubated for 1 h at room temperature with secondary HRP-conjugated goat anti-mouse goat antibodies (Invitrogen #31430, 1:5000 dilution). The blot was developed using SuperSignal West Femto Maximum Sensitivity ECL Substrate (Thermo Fischer Scientific 34094) on a VersaDoc Imaging System (Bio-Rad 4000 MP).

### Loci tracking and trajectory analysis

The localization of each fluorescently-labeled chromatin focus was determined by applying a spatial bandpass filter to each raw image frame. Specifically, the centroid of each peak above a determined threshold value was calculated to obtain an approximate position. Then, the final localization was obtained by fitting a radially symmetric Gaussian function around each centroid position. These positions were then linked into time trajectories using the algorithm for single particle tracking introduced in (Crocker and Grier, 1996). Specifically, we did not allow gaps between frames, we set the maximum displacement between frames to 2 pixels (0.321 μm), and we set the minimum track length to be 300 frames, or 100 frames for experiments involving *arp8Δ*, *arp9Δ*, and *cut14-208/arp8Δ* genotypes or sodium azide treatment. The mean-square displacement (MSD) versus time delay for each time-lapse image series was calculated as follows. First, the trajectories were split into sub-trajectories of 29 steps (with duration ≈ 1.68 s).

Then, individual displacements were averaged across all sub-trajectories and different time-lapse series for a given experiment. The calculated experimental MSDs were fit by the theoretical MSD for fractional Brownian motion (fBm) with static and dynamic localization errors taken into account, as described (Bailey et al., 2021):

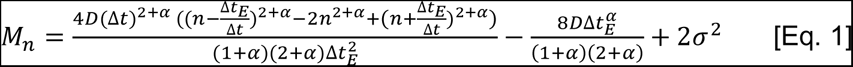

where *D* is an effective diffusivity, Δ*t*_*E*_ = 10 ms is exposure time, Δ*t* is the time step between frames, and α is an exponent (α < 1 corresponds to sub-diffusive motion and α > 1 to super-diffusive motion), *n* is the number of frames, and *σ* is the static localization error. Initial MSD fitting, in which *D*, α, and *σ* were varied, showed that the values of α grouped around a mean of 0.44 for most measured loci in unperturbed, wild type cells (Supplemental Table S1). In all subsequent analyses (see Figs. 1, 2b, 3, 5, 7), α was fixed at α = 0.44, while *D* and *σ* were varied, allowing us to directly compare values of *D* as a measure of locus mobility across different experiments. Supplemental Table S1 contains all the fitting results.

### Whole-nucleus motion tracking

To estimate the potential contribution of rigid body motion of the nucleus to our measurements of chromatin dynamics, we analyzed the dynamics of nuclei in a strain expressing a fluorescently-tagged form of the nucleoporin, Cut11 (Cut11-mCherry). Single central plane fluorescent images were acquired every 2 seconds. First, approximate nuclei positions were detected using a custom-written routine in MATLAB. Next, each detected fluorescent image of the nuclear envelope rim was fitted (using custom-written routine in MATLAB) by a circle convolved with the point-spread function with the center position, circled radius, fluorescence amplitude and background as free parameters. Last, the position of each nuclear center was tracked by the same approach described for loci tracking and trajectory analysis.

### Loop-extrusion-factor (LEF) simulations

We carried out Gillespie-type simulations (Gillespie, 1977) of the chromatin loop configuration using two different models of loop formation and extrusion. First, we simulated a version of the LEF model described in (Fudenberg et al., 2016). Subsequent to uniformly-distributed random binding, LEF translocation proceeds stochastically and bidirectionally at a certain rate *v* until either the LEF dissociates or a LEF anchor encounters another LEF or a boundary element (BE). Only outward translocation (i.e. one increasing the loop) is permitted, with translocation at each loop anchor occurring randomly and independently with equal probability. LEFs cannot pass each other; however, boundary elements (BEs) have a non-zero permeability, *P*, which is implemented as a multiplier to the LEF translocation rate at BEs, that is, the translocation rate through BEs *v_BE_* = *P v*. CTCF ChIP-seq data (Bonev et al., 2017) was used to determine the locations of boundary element (BEs), as well as their permeability *P*, which was determined from the CTCF ChIP-seq signal *S* via a logistic function:

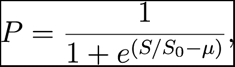

where *S*_0_ and *μ* are free parameters to choose. We set *S*_0_ = 20 and *μ* = 3 following the parameters used in (Fudenberg *et al*., 2016). Second, since *S. pombe* lack CTCF, we also carried out simulations without any BEs, called the “LEF only” model, which is otherwise the same as the previous model. For each set of parameter values, we performed 30 simulations, each comprising 100,000 time steps. Loop dynamics reached a steady state over about 5,000 steps as gauged by convergence of the chromatin backbone length, as well as the chromatin’s radius of gyration, to a more-or-less constant value. 6 Mb (300 kb) genomic regions for mouse (yeast) were divided into 10 kb (0.5 kb) bins, creating 600 sites that could be occupied by an anchor of a LEF. The LEF model parameters used for both CTCF and random loop models for both mouse and yeast are presented in Supplemental Table S2. The parameters used closely follow from the range of values given in (Fudenberg et al., 2016). For simulations considering two type of LEFs, an equal number of LEFs with two different dissociation times that bind and move independently of each other (unless they encounter another LEF of the same or different type) were simulated. Our LEF simulation code is online at the Github repository: https://github.com/nilesyan/LEF-Simulation.

### Rouse-model polymer simulations

Chromatin’s polymer behavior was modeled as a free-end Rouse (bead-spring) chain, in which each 10 kb (0.5 kb) region of the mouse (yeast) genome is represented as a bead that is connected to its nearest neighbors by springs of stiffness *κ*, and experiences fluid friction with coefficient *ζ*. By treating each Rouse segment as an entropic spring, *κ* is derived from the chromatin persistence length *l*_*k*_ estimated in (Arbona et al., 2017) using the formula 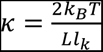, where *L* is the contour length of the segment. Given the calculated *κ* and measured diffusion coefficient *D* from our experiments, the friction coefficient *ζ* can be evaluated by the formula 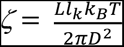, which is derived from classical Rouse model (Doi and Edwards, 1986). For the simulations shown in Fig. 4-6, the characteristic polymer time, defined here as the reciprocal of the largest Rouse-model eigenvalue, namely 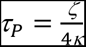, was then calculated and used as a time unit (Supplemental Table S2). To include loops, we augmented the model with additional springs (of the same spring constant) that can connect non-adjacent beads, representing the base of a loop. The sizes and locations of loops evolve according to the loop configurations generated by the LEF model simulation described above. Since the additional springs (loops) lead to far-from-diagonal terms in the Rouse-model dynamical matrix, the matrix was diagonalized numerically to find the eigenmodes and corresponding beads-to-eigenmode transformations. Assuming equipartition with an effective temperature, these eigenmodes (and, through transformations, the corresponding bead positions) were then evolved using the exact simulation method described in (Gillespie, 1996) until the loop configuration changed. This procedure was then repeated for each new loop configuration. At any timepoint when the loop configuration changes, we set the beads’ positions immediately after the timepoint to be the same as the beads’ positions immediately before the timepoint. Therefore, the beads’ positions evolve continuously, and we can simulate the motion of the chromatin beads in an exact and continuous manner. As a result, we are able to simulate polymer dynamics exactly with arbitrary time steps in our simulations and combine an equally-spaced time series (with time step comparable to the polymer time) with the exact times of loop-extrusion events, ensuring a recapitulation of the chromatin motion in the presence of loop-extrusion. Furthermore, since the polymer relaxation time is much shorter than the characteristic loop-extrusion time, the loop dynamics due to active loop-extrusion has a negligible effect on the chromatin polymer dynamics. Finally, for the simulations shown in Fig. 4-6, we generated an independent Rouse polymer simulation for each of 30 LEF simulations and chose 60 equally-separated beads along the chromatin polymer. Each bead’s MSD and effective exponent results were averaged over these 30 independent polymer simulations (shown in light colors), and the dark colored lines are the averaged results of the 60 individual beads (light colored lines).

## Contributions

J.F.W., K.L., K.D. and I.S. generated all yeast strains and raw movies for analysis of chromatin mobility. M.P.B. designed, implemented and plotted all loci tracking and trajectory analysis. H.Y. and T.Y. designed and performed the Gillespie and polymer simulations with input from I.S. and S.G.J.M. M.C.K. and S.G.J.M. conceived of the project, coordinated the team strategy and provided experimental direction. I.S. led the integration of the experimental and simulation approaches. The manuscript was written, edited, and approved by all authors.

## Acknowledgements

We thank Bryan Leland, Amanda Vines, Paolo Colombi, Dongxu Lin, Sara Siwiecki, and Ryan Brennan for generating strains used in this work. We also thank the Yeast Genome Resource Center at Osaka University and the many investigators who have deposited strains at this resource. This work was supported by an NSF EFRI CEE award EFMA-1830904 to S.G.J. Mochrie and M.C. King. M.P.B. was supported by NIH T32EB019941 and the NSF GRFP. H.Y. was supported by the NSF Physics of Living Systems Student Research Network via NSF PHYS 1522467.

---

**Supplemental Table S1:**
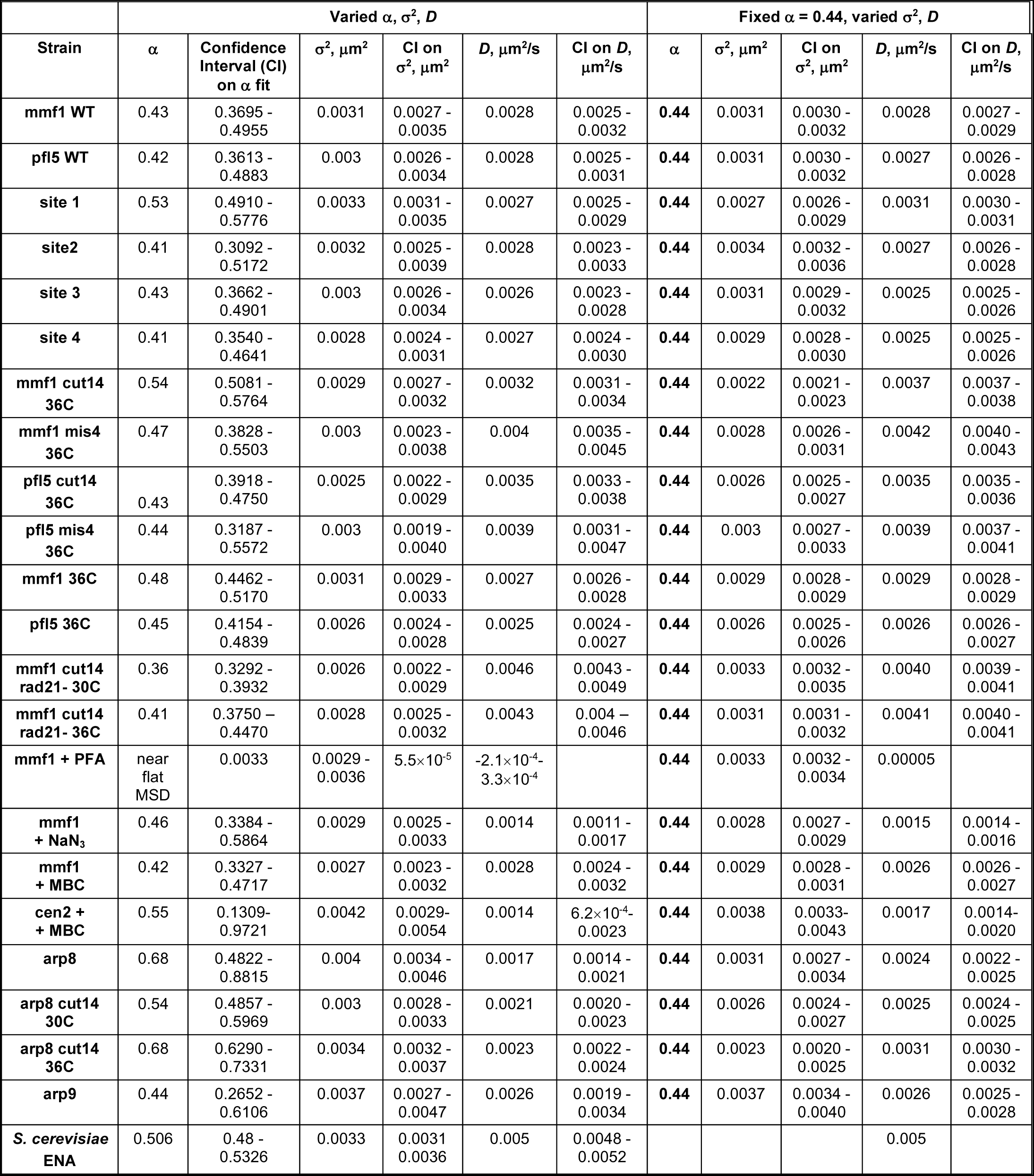
MSD Fitting Results

**Supplemental Table S2:**
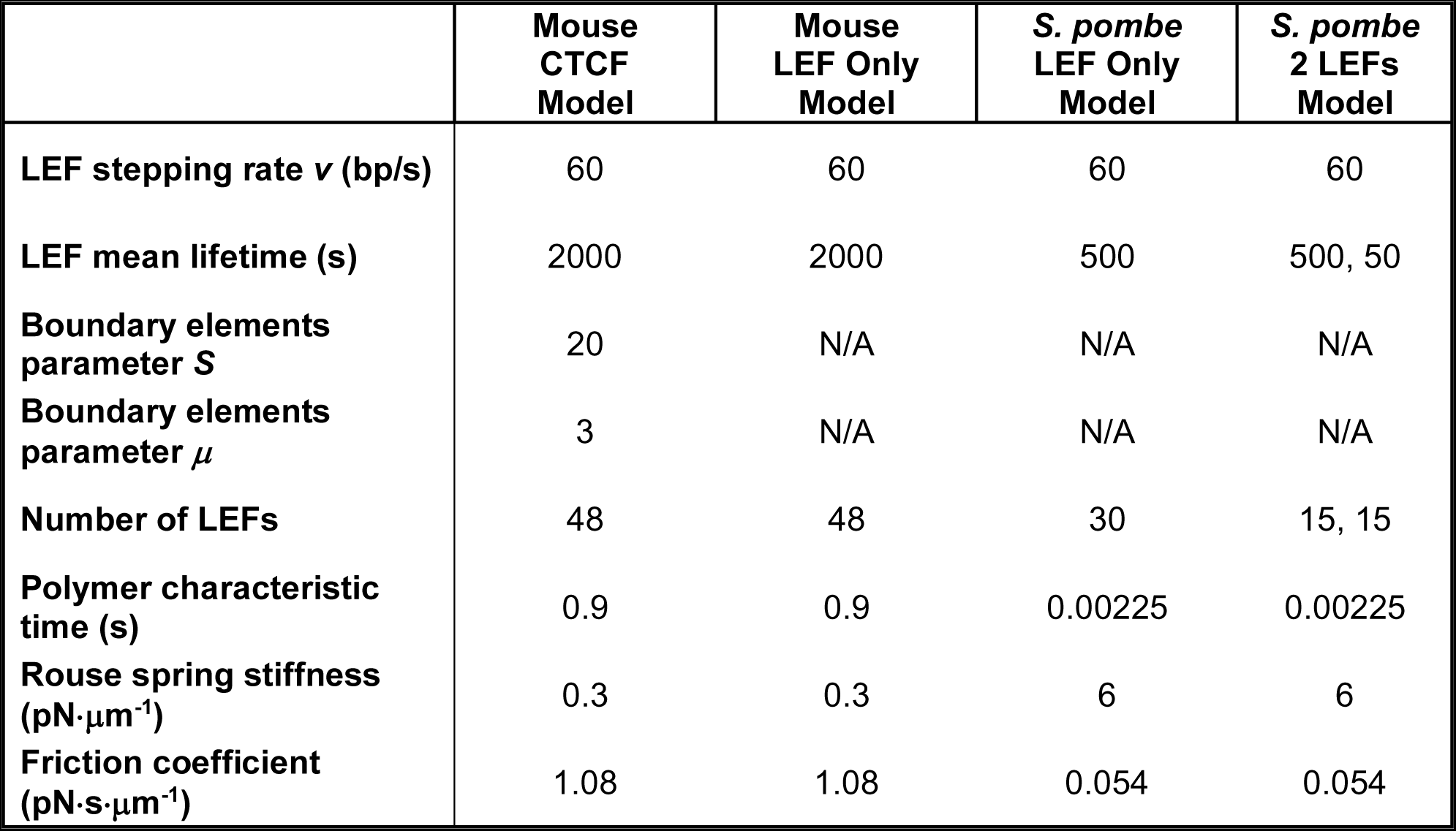
Loop-extrusion-factor (LEF) Simulation and Rouse Simulation Parameters

|  | Mouse<br>CTCF<br>Model | Mouse<br>LEF Only<br>Model | *S. pombe*<br>LEF Only<br>Model | *S. pombe*<br>2 LEFs<br>Model |
| --- | --- | --- | --- | --- |
| LEF stepping rate $v$ (bp/s) | 60 | 60 | 60 | 60 |
| LEF mean lifetime (s) | 2000 | 2000 | 500 | 500, 50 |
| Boundary elements<br>parameter $S$ | 20 | N/A | N/A | N/A |
| Boundary elements<br>parameter $\mu$ | 3 | N/A | N/A | N/A |
| Number of LEFs | 48 | 48 | 30 | 15, 15 |
| Polymer characteristic<br>time (s) | 0.9 | 0.9 | 0.00225 | 0.00225 |
| Rouse spring stiffness<br>($\text{pN}\cdot\mu\text{m}^{-1}$) | 0.3 | 0.3 | 6 | 6 |
| Friction coefficient<br>($\text{pN}\cdot\text{s}\cdot\mu\text{m}^{-1}$) | 1.08 | 1.08 | 0.054 | 0.054 |

**Supplemental Table S3.** Strains used in the study

| Strain | Relevant genotype | Description | Source |
| --- | --- | --- | --- |
| ***S. pombe*** |  |  |  |
| *Specific lacO position* |  |  |  |
| MKSP1642 | *h+ leu1-32 ura4-D18*<br>*his7+::P<sub>Dis1</sub>-gfp-lacI-NLS*<br>*sad1::sad1-dsRed-leu+*<br>*cen2::ura4-lacO<sub>n</sub>* | WT strain with *lacO* array integrated near *cen2* (Chr2 centromere) locus and with expression of LacI-GFP (with nucleus localization signal) under control of the constitutive promoter. *Fig. 2.* | *this study derived from PX342 and PN10127* |
| MKSP1661 | *h- leu1-32 ura4-D18*<br>*his7+::P<sub>Dis1</sub>-gfp-lacI-NLS*<br>*cut11::cut11-mCherry-natR*<br>*pfl5::ura4-lacO<sub>n</sub>* | WT strain with *lacO* array (5.6 kb) integrated near *pfl5* locus (at *ChrII*: 4403547-4403553). *Fig. 1b, 3b, Fig. S1a.* | [1] |
| MKSP1794 | *leu1-32 ura4-D18*<br>*mis4::mis4-242<sup>ts</sup>*<br>*his7+::P<sub>Dis1</sub>-gfp-lacI-NLS*<br>*cut11::cut11-mCherry-natR*<br>*pfl5::ura4-lacO<sub>n</sub>* | Temperature-sensitive mutant of Mis4 cohesin loading factor with *lacO* array (5.6 kb) integrated near *pfl5* locus. *Fig. 3b.* | *this study derived from MY3655 and MKSP1340* |
| MKSP2039 | *h+ leu1-32 ura4-D18*<br>*his7+::P<sub>Dis1</sub>-gfp-lacI-NLS*<br>*mmf1::ura4-lacO<sub>n</sub>* | WT strain with *lacO* array (10.3 kb) integrated near *mmf1* locus (at *ChrII*:3442981). *Fig. 1b-c, 2, 3a, 5, 7, Fig. S1a, S2* | *this study derived from MKSP1381* |
| MKSP2583 | *h- leu1-32 ura4-D18*<br>*arp8::hygR*<br>*his7+::P<sub>Dis1</sub>-gfp-lacI-NLS*<br>*mmf1::ura4-lacO<sub>n</sub>*<br>*urg1::loxP-kanR-loxM3* | *arp8* (INO80 complex subunit) deletion mutant with *lacO* array (10.3 kb) integrated near *mmf1* locus. *Fig. 7.* | *this study derived from MKSP2190* |
| MKSP2693 | *leu1-32 ura4-D18*<br>*cut14::cut14-208<sup>ts</sup>*<br>*cut11::cut11-mCherry-natR*<br>*his7+::P<sub>Dis1</sub>-gfp-lacI-NLS*<br>*pfl5::ura4-lacO<sub>n</sub>* | Temperature-sensitive mutant of Cut14 condensin subunit with *lacO* array (5.6 kb) integrated near *pfl5* locus. *Fig. 3b.* | *this study derived from MY1983* |
| MKSP2760 | *leu1-32 ura4-D18*<br>*cut14::cut14-208<sup>ts</sup>*<br>*his7+::P<sub>Dis1</sub>-gfp-lacI-NLS*<br>*mmf1::ura4-lacO<sub>n</sub>* | Temperature-sensitive mutant of Cut14 condensin subunit with *lacO* array (10.3 kb) integrated near *mmf1* locus. *Fig. 3a, 5b.* | *this study derived from MY1997* |
| MKSP2801 | *leu1-32 ura4-D18*<br>*mis4::mis4-242<sup>ts</sup>*<br>*his7+::P<sub>Dis1</sub>-gfp-lacI-NLS*<br>*cut11::cut11-mCherry-natR*<br>*mmf1::ura4-lacO<sub>n</sub>* | Temperature-sensitive mutant of Mis4 cohesin loading factor with *lacO* array (10.3 kb) integrated near *mmf1* locus. *Fig. 3a.* | *this study derived from MY3655 and MKSP1340* |
| MKSP3021 | *leu1-32 ura4-D18*<br>*arp9::kanR*<br>*his7+::P<sub>Dis1</sub>-gfp-lacI-NLS*<br>*mmf1::ura4-lacO<sub>n</sub>* | *arp9* (SWI/RSC complexes subunit) deletion with *lacO* array (10.3 kb) integrated near *mmf1* locus. *Fig. 7a.* | *this study derived from MKSP2352* |
| MKSP3053 | *leu1-32 ura4-D18*<br>*arp8::kanR cut14::cut14-208<sup>ts</sup>*<br>*his7+::P<sub>Dis1</sub>-gfp-lacI-NLS*<br>*mmf1::ura4-lacO<sub>n</sub>* | Double mutant with *arp8* deletion and temperature-sensitive mutant of Cut14 condensin subunit with *lacO* array (10.3 kb) integrated near *mmf1* locus. *Fig. 7b.* | *this study derived from MKSP2187* |
| MKSP3660 | *leu1-32 cut14::cut14-208<sup>ts</sup> his7+:P<sub>Dis1</sub>-gfp-lacI-NLS mmf1::ura4-lacO<sub>n</sub> rad21::rad21-XTEN17-3sAID-kanR ura4::ura4-P<sub>adh1</sub>-OsTir1-F74A* | Double mutant bearing temperature-sensitive allele of Cut14 condensin subunit and rad21 cohesin subunit tagged with auxin-inducible degron with *lacO* array (10.3 kb) integrated near *mmf1* locus. Fig. 5b | *this study derived from DY48569* |
| *Random lacO position* |  |  |  |
| MKSP3117 | *h+ leu1-32 ura4-D18 his7+:P<sub>Dis1</sub>-gfp-lacI-NLS random locus:ura4-lacO<sub>n</sub>* | WT strain with *lacO* array (10.3 kb) integrated at random locus of the genome (noted as "site1"), and with LacI-GFP (with nucleus localization signal) under control of the constitutive promoter. Fig. 1b, Fig. S1a. | *derived from MKSP1120* |
| MKSP3118 | *h+ leu1-32 ura4-D18 his7+:P<sub>Dis1</sub>-gfp-lacI-NLS random locus:ura4-lacO<sub>n</sub>* | Same as MKSP3117 but with a different random *lacO* array position (noted as "site2"). Fig. 1b, Fig. S1a. | *derived from MKSP1120* |
| MKSP3119 | *h+ leu1-32 ura4-D18 his7+:P<sub>Dis1</sub>-gfp-lacI-NLS random locus:ura4-lacO<sub>n</sub>* | Same as MKSP3117 but with a different random *lacO* array position (noted as "site3"). Fig. 1b, Fig. S1a. | *derived from MKSP1120* |
| MKSP3123 | *h+ leu1-32 ura4-D18 his7+:P<sub>Dis1</sub>-gfp-lacI-NLS random locus:ura4-lacO<sub>n</sub>* | Same as MKSP3117 but with a different random *lacO* array position (noted as "site4"). Fig. 1b, Fig. S1a. | *derived from MKSP1120* |
| *Labeled nuclear envelope strain* |  |  |  |
| MKSP3140 | *h+ leu1-32 ura4-D18 his7+:P<sub>Dis1</sub>-gfp-lacI-NLS cut11::cut11-mCherry-natR isd90: ura4-lacO<sub>n</sub>* | WT strain with *lacO* array (10 kb) integrated near *isd90* locus (at *ChrII*: 1950422). Fig. S2. |  |
| *Auxiliary strains* |  |  |  |
| AW563 | *h- leu1-32 urg1:loxP-kanR-loxM3<sub>6</sub>* | WT strain with recombination-mediated cassette exchange system (*RMCE<sub>kanMX6</sub>*). | [2] |
| DY48569 | *h+ mat1PΔ17 leu1-32 lys1-131 ade6-M216 ura4::ura4-P<sub>adh1</sub>-OsTir1-F74A* | Strain bearing an auxin-inducible degron system | YGRC FY39923 [3] |
| JM210<br>FN198<br>MKSP1116 | *h- leu1-32 ura4-D18 ade-cut11::cut11-mCherry-natMX6* | WT strain with Cut11-mCherry (a nuclear envelope marker) expression controlled by the native promoter. | *Nurse lab* |
| MKSP1120 | *h+ leu1-32 ura4-D18 his7+:P<sub>Dis1</sub>-gfp-lacI-NLS* | WT strain with LacI-GFP (with nucleus localization signal) under control of the constitutive promoter. Used to generate random *lacO* array integrations. |  |
| MKSP1340 | *h- leu1-32 ura4-D18 his7+:P<sub>Dis1</sub>-gfp-lacI-NLS cut11::cut11-mCherry-natR mmf1::ura4-lacO<sub>n</sub> ChrII:3442981::ura4-lacO<sub>n</sub>* | WT strain with *lacO* array (10.3 kb) integrated near *mmf1* locus and with LacI-GFP (with nucleus localization signal) under control of the constitutive promoter | *this study similar to MKSP1381* |
| MKSP1381 | *h- leu1-32 ura4-D18 his7+:P<sub>Dis1</sub>-gfp-lacI-NLS ChrII:3442981::ura4-10.3kbLacO ChrII:3446249::HOcs-hphMX6 cut11::cut11-mCherry-natMX6* | Original *lacO* array (10.3 kb) integration near *mmf1* locus. Used to generate all other *mmf1-lacO<sub>n</sub>* strains in this study | [4] |
| MKSP1128 | *h- leu1-32 ura4-D18 his7+:P<sub>Dis1</sub>-gfp-lacI-NLS cut11::cut11-mCherry-natMX6* | WT strain with Cut11-mCherry (a nuclear envelope marker) expression controlled by the native promoter and with LacI-GFP (with nucleus localization signal) under control of the constitutive promoter. |  |
| MKSP2187 | *h- leu1-32 ura4-D18 arp8::kanR* | *arp8* deletion. | *this study* |
| MKSP2190 | *h- leu1-32 ura4-D18 arp8::hygR* | *arp8* deletion. | *this study* |
| MKSP2352 | *h- leu1-32 ura4-D18 arp9::kanR* | *arp9* deletion. | *this study* |
| MY1983 | *h- ura4 cut14::cut14-208<sup>ts</sup>* | Strain encoding for the temperature sensitive allele of *cut14*. | YGRC FY9883 |
| MY1997 | *h+ cut14::cut14-208<sup>ts</sup>* | Strain encoding for the temperature sensitive allele of *cut14*. | YGRC FY9897 |
| MY3655 | *h- leu1-32 mis4::mis4-242<sup>ts</sup>* | Strain encoding for the temperature sensitive allele of *mis4*. | YGRC FY111578 |
| PN10127 | *h- sad1::sad1-dsRed-leu+* | WT strain with Sad1-dsRed (a spindle pole body marker) expression controlled by the native promoter. |  |
| PX342 | *h+ leu1-32 ade6-M216 bub1::ura4 cen2-lacO<sub>n</sub>* | Strain with *cen2* locus (centromere of ChrII) containing *lacO* array insertion. | YGRC FY13812 [5] |
| ***S. cerevisiae*** |  |  |  |
| PCCPL645 | *W303 LacI-GFP::HIS* | W303 strain with *LacI-GFP* (with nucleus localization signal) under control of the constitutive promoter. | [6] |
| PCCPL835 | *W303 LacI-GFP::HIS ENA-lacO<sub>n</sub>::TRP1* | W303 strain with *lacO* array integrated near *ENA* locus and with *LacI-GFP* (with nucleus localization signal) under control of the constitutive promoter. *Fig. S1b*. | *this study derived from PCCPL 645 as described [6] and [7]* |

1. Zhao Y., Schreiner S.M., Koo P.K., Colombi P., King M.C., Mochrie S.G. (2016) Improved Determination of Subnuclear Position Enabled by Three-Dimensional Membrane Reconstruction. **Biophys. J.** 111, 19-24. 2. Watson A.T., Werler P., Carr A.M. (2011) Regulation of gene expression at the fission yeast *Schizosaccharomyces pombe* *urg1* locus. **Gene** 484, 75-85. 3. Zhang X.-R. Zhao L., Suo F., Gao Y., Wu Q., Qi X., Du L.-L. (2022) An improved auxin-inducible degron system for fission yeast. **G3**, 12, jkab393. 3. Leland B.A., Chen A.C., Zhao A.Y., Wharton R.C., King M.C. (2018) Rev7 and 53BP1/Crb2 prevent RecQ helicase-dependent hyper-resection of DNA double-strand breaks. **eLife** 7, e33402. 4. Hauf S., Biswas A., Langeegger M., Kawashima S.A., Tsukahara T., Watanabe Y. (2007) Aurora controls sister kinetochore mono-orientation and homolog bi-orientation in meiosis-I. **EMBO J.** 26, 4475-86. 5. Colombi P., King D.E., Williams J.F., Lusk C.L., King M.C. (2018) LEM domain proteins control the efficiency of adaptation through copy number variation. **Bioarxiv** 451583, https://www.biorxiv.org/content/biorxiv/early/2018/10/24/451583.full.pdf 6. Murray J., Watson A., Carr A. (2016) *Identifying products of recombinase-mediated cassette exchange (RMCE) in Schizosaccharomyces pombe*. **Cold Spring Harbor Protocols**, 2016 (5)

## References

Alipour, E., and J.F. Marko. 2012. Self-organization of domain structures by DNA-loop-extruding enzymes. Nucleic acids research. 40:11202–11212.

Arbona, J.M., S. Herbert, E. Fabre, and C. Zimmer. 2017. Inferring the physical properties of yeast chromatin through Bayesian analysis of whole nucleus simulations. Genome Biol. 18:81.

Bähler, J., J.Q. Wu, M.S. Longtine, N.G. Shah, A. McKenzie, A.B. Steever, A. Wach, P. Philippsen, and J.R. Pringle. 1998. Heterologous modules for efficient and versatile PCR-based gene targeting in Schizosaccharomyces pombe. *Yeast (Chichester*, England). 14:943–951.

Bailey, M.L.P., H. Yan, I. Surovtsev, J.F. Williams, M.C. King, and S.G.J. Mochrie. 2021. Covariance distributions in single particle tracking. Physical Review E. 103:032405.

Banigan, E.J., and L.A. Mirny. 2020. Loop extrusion: theory meets single-molecule experiments. Current opinion in cell biology. 64:124–138.

Banigan, E.J., A.A. van den Berg, H.B. Brandão, J.F. Marko, and L.A. Mirny. 2020. Chromosome organization by one-sided and two-sided loop extrusion. Elife. 9.

Batut, P.J., X.Y. Bing, Z. Sisco, J. Raimundo, M. Levo, and M.S. Levine. 2022. Genome organization controls transcriptional dynamics during development. *Science (New York*, N.Y.). 375:566–570.

Bintu, B., L.J. Mateo, J.H. Su, N.A. Sinnott-Armstrong, M. Parker, S. Kinrot, K. Yamaya, A.N. Boettiger, and X. Zhuang. 2018. Super-resolution chromatin tracing reveals domains and cooperative interactions in single cells. *Science (New York*, N.Y.). 362.

Bonev, B., N. Mendelson Cohen, Q. Szabo, L. Fritsch, G.L. Papadopoulos, Y. Lubling, X. Xu, X. Lv, J.P. Hugnot, A. Tanay, and G. Cavalli. 2017. Multiscale 3D Genome Rewiring during Mouse Neural Development. Cell. 171:557–572 e524.

Caridi, C.P., C. D’Agostino, T. Ryu, G. Zapotoczny, L. Delabaere, X. Li, V.Y. Khodaverdian, N. Amaral, E. Lin, A.R. Rau, and I. Chiolo. 2018. Nuclear F-actin and myosins drive relocalization of heterochromatic breaks. Nature. 559:54–60.

Cheblal, A., K. Challa, A. Seeber, K. Shimada, H. Yoshida, H.C. Ferreira, A. Amitai, and S.M. Gasser. 2020. DNA Damage-Induced Nucleosome Depletion Enhances Homology Search Independently of Local Break Movement. Mol Cell. 80:311–326.e314.

Chuang, C.H., A.E. Carpenter, B. Fuchsova, T. Johnson, P. de Lanerolle, and A.S. Belmont. 2006. Long-range directional movement of an interphase chromosome site. Curr Biol. 16:825–831.

Colombi, P., D.E. King, J.F. Williams, C.P. Lusk, and M.C. King. 2018. LEM domain proteins control the efficiency of adaptation through copy number variation. bioRxiv:451583.

Crocker, J.C., and D.G. Grier. 1996. Methods of Digital Video Microscopy for Colloidal Studies. Journal of Colloid and Interface Science. 179:298–310.

D’Ambrosio, C., C.K. Schmidt, Y. Katou, G. Kelly, T. Itoh, K. Shirahige, and F. Uhlmann. 2008. Identification of cis-acting sites for condensin loading onto budding yeast chromosomes. Genes & development. 22:2215–2227.

Davidson, I.F., B. Bauer, D. Goetz, W. Tang, G. Wutz, and J.M. Peters. 2019. DNA loop extrusion by human cohesin. *Science (New York*, N.Y.). 366:1338–1345.

Dekker, J. 2014. Two ways to fold the genome during the cell cycle: insights obtained with chromosome conformation capture. Epigenetics Chromatin. 7:25.

Dion, V., V. Kalck, C. Horigome, B.D. Towbin, and S.M. Gasser. 2012. Increased mobility of double-strand breaks requires Mec1, Rad9 and the homologous recombination machinery. Nature cell biology. 14:502-509.

Dixon, J.R., D.U. Gorkin, and B. Ren. 2016. Chromatin Domains: The Unit of Chromosome Organization. Mol Cell. 62:668–680.

Dixon, J.R., S. Selvaraj, F. Yue, A. Kim, Y. Li, Y. Shen, M. Hu, J.S. Liu, and B. Ren. 2012. Topological domains in mammalian genomes identified by analysis of chromatin interactions. Nature. 485:376–380.

Doi, M., and S.F. Edwards. 1986. The theory of polymer dynamics. Oxford University Press, New York.

Dubey, R.N., and M.R. Gartenberg. 2007. A tDNA establishes cohesion of a neighboring silent chromatin domain. Genes & development. 21:2150–2160.

Falk, M., Y. Feodorova, N. Naumova, M. Imakaev, B.R. Lajoie, H. Leonhardt, B. Joffe, J. Dekker, G. Fudenberg, I. Solovei, and L.A. Mirny. 2019. Heterochromatin drives compartmentalization of inverted and conventional nuclei. Nature. 570:395–399.

Flavahan, W.A., Y. Drier, B.B. Liau, S.M. Gillespie, A.S. Venteicher, A.O. Stemmer-Rachamimov, M.L. Suva, and B.E. Bernstein. 2016. Insulator dysfunction and oncogene activation in IDH mutant gliomas. Nature. 529:110–114.

Fudenberg, G., M. Imakaev, C. Lu, A. Goloborodko, N. Abdennur, and L.A. Mirny. 2016. Formation of Chromosomal Domains by Loop Extrusion. Cell reports. 15:2038–2049.

Funabiki, H., I. Hagan, S. Uzawa, and M. Yanagida. 1993. Cell cycle-dependent specific positioning and clustering of centromeres and telomeres in fission yeast. J Cell Biol. 121:961–976.

Gabriele, M., H.B. Brandão, S. Grosse-Holz, A. Jha, G.M. Dailey, C. Cattoglio, T.S. Hsieh, L. Mirny, C. Zechner, and A.S. Hansen. 2022. Dynamics of CTCF- and cohesin-mediated chromatin looping revealed by live-cell imaging. *Science (New York*, N.Y.). 376:496–501.

Ganji, M., I.A. Shaltiel, S. Bisht, E. Kim, A. Kalichava, C.H. Haering, and C. Dekker. 2018. Real-time imaging of DNA loop extrusion by condensin. *Science (New York*, N.Y.). 360:102–105.

Garcia-Luis, J., L. Lazar-Stefanita, P. Gutierrez-Escribano, A. Thierry, A. Cournac, A. Garcia, S. Gonzalez, M. Sanchez, A. Jarmuz, A. Montoya, M. Dore, H. Kramer, M.M. Karimi, F. Antequera, R. Koszul, and L. Aragon. 2019. FACT mediates cohesin function on chromatin. Nature structural & molecular biology. 26:970–979.

Gassler, J., H.B. Brandao, M. Imakaev, I.M. Flyamer, S. Ladstatter, W.A. Bickmore, J.M. Peters, L.A. Mirny, and K. Tachibana. 2017. A mechanism of cohesin-dependent loop extrusion organizes zygotic genome architecture. EMBO J. 36:3600–3618.

Gerlich, D., T. Hirota, B. Koch, J.M. Peters, and J. Ellenberg. 2006a. Condensin I stabilizes chromosomes mechanically through a dynamic interaction in live cells. Curr Biol. 16:333–344.

Gerlich, D., B. Koch, F. Dupeux, J.M. Peters, and J. Ellenberg. 2006b. Live-cell imaging reveals a stable cohesin-chromatin interaction after but not before DNA replication. Curr Biol. 16:1571–1578.

Gillespie, D.T. 1977. Exact stochastic simulation of coupled chemical reactions. The Journal of Physical Chemistry. 81:2340–2361.

Gillespie, D.T. 1996. The mathematics of Brownian motion and Johnson noise. Am J Phys.64:225–240.

Glynn, E.F., P.C. Megee, H.G. Yu, C. Mistrot, E. Unal, D.E. Koshland, J.L. DeRisi, and J.L. Gerton. 2004. Genome-wide mapping of the cohesin complex in the yeast Saccharomyces cerevisiae. PLoS Biol. 2:E259.

Golfier, S., T. Quail, H. Kimura, and J. Brugués. 2020. Cohesin and condensin extrude DNA loops in a cell cycle-dependent manner. Elife. 9.

Gu, B., T. Swigut, A. Spencley, M.R. Bauer, M. Chung, T. Meyer, and J. Wysocka. 2018. Transcription-coupled changes in nuclear mobility of mammalian cis-regulatory elements. *Science (New York*, N.Y.). 359:1050–1055.

Haarhuis, J.H.I., R.H. van der Weide, V.A. Blomen, J.O. Yanez-Cuna, M. Amendola, M.S. van Ruiten, P.H.L. Krijger, H. Teunissen, R.H. Medema, B. van Steensel, T.R. Brummelkamp, E. de Wit, and B.D. Rowland. 2017. The Cohesin Release Factor WAPL Restricts Chromatin Loop Extension. Cell. 169:693–707 e614.

Haering, C.H., A.M. Farcas, P. Arumugam, J. Metson, and K. Nasmyth. 2008. The cohesin ring concatenates sister DNA molecules. Nature. 454:297–301.

Hauer, M.H., A. Seeber, V. Singh, R. Thierry, R. Sack, A. Amitai, M. Kryzhanovska, J. Eglinger, D. Holcman, T. Owen-Hughes, and S.M. Gasser. 2017. Histone degradation in response to DNA damage enhances chromatin dynamics and recombination rates. Nature structural & molecular biology. 24:99–107.

Hentges, P., B. Van Driessche, L. Tafforeau, J. Vandenhaute, and A.M. Carr. 2005. Three novel antibiotic marker cassettes for gene disruption and marker switching in Schizosaccharomyces pombe. Yeast. 22:1013–1019.

Heun, P., T. Laroche, M.K. Raghuraman, and S.M. Gasser. 2001. The positioning and dynamics of origins of replication in the budding yeast nucleus. J Cell Biol. 152:385–400.

Hirano, T. 2016. Condensin-Based Chromosome Organization from Bacteria to Vertebrates. Cell. 164:847–857.

Hu, B., T. Itoh, A. Mishra, Y. Katoh, K.L. Chan, W. Upcher, C. Godlee, M.B. Roig, K. Shirahige, and K. Nasmyth. 2011. ATP hydrolysis is required for relocating cohesin from sites occupied by its Scc2/4 loading complex. Curr Biol. 21:12–24.

Ibrahim, D.M., and S. Mundlos. 2020. The role of 3D chromatin domains in gene regulation: a multi-facetted view on genome organization. Current opinion in genetics & development. 61:1–8.

Jo, H., T. Kim, Y. Chun, I. Jung, and D. Lee. 2021. A compendium of chromatin contact maps reflecting regulation by chromatin remodelers in budding yeast. Nature communications. 12:6380.

Joyner, R.P., J.H. Tang, J. Helenius, E. Dultz, C. Brune, L.J. Holt, S. Huet, D.J. Müller, and K. Weis. 2016. A glucose-starvation response regulates the diffusion of macromolecules. Elife. 5.

Kakui, Y., C. Barrington, D.J. Barry, T. Gerguri, X. Fu, P.A. Bates, B.S. Khatri, and F. Uhlmann. 2020. Fission yeast condensin contributes to interphase chromatin organization and prevents transcription-coupled DNA damage. Genome Biol. 21:272.

Kim, Y., Z. Shi, H. Zhang, I.J. Finkelstein, and H. Yu. 2019. Human cohesin compacts DNA by loop extrusion. *Science (New York*, N.Y.). 366:1345–1349.

Lawrimore, C.J., J. Lawrimore, Y. He, S. Chavez, and K. Bloom. 2020. Polymer perspective of genome mobilization. Mutation research. 821:111706.

Leland, B.A., and M.C. King. 2014. Using LacO arrays to monitor DNA double-strand break dynamics in live Schizosaccharomyces pombe cells. *Methods in molecular biology (Clifton*, N.J.). 1176:127–141.

Lengronne, A., Y. Katou, S. Mori, S. Yokobayashi, G.P. Kelly, T. Itoh, Y. Watanabe, K. Shirahige, and F. Uhlmann. 2004. Cohesin relocation from sites of chromosomal loading to places of convergent transcription. Nature. 430:573–578.

Lieberman-Aiden, E., N.L. van Berkum, L. Williams, M. Imakaev, T. Ragoczy, A. Telling, I. Amit, B.R. Lajoie, P.J. Sabo, M.O. Dorschner, R. Sandstrom, B. Bernstein, M.A. Bender, M. Groudine, A. Gnirke, J. Stamatoyannopoulos, L.A. Mirny, E.S. Lander, and J. Dekker. 2009. Comprehensive mapping of long-range interactions reveals folding principles of the human genome. *Science (New York*, N.Y.). 326:289–293.

Lottersberger, F., R.A. Karssemeijer, N. Dimitrova, and T. de Lange. 2015. 53BP1 and the LINC Complex Promote Microtubule-Dependent DSB Mobility and DNA Repair. Cell. 163:880–893.

Lupianez, D.G., K. Kraft, V. Heinrich, P. Krawitz, F. Brancati, E. Klopocki, D. Horn, H. Kayserili, J.M. Opitz, R. Laxova, F. Santos-Simarro, B. Gilbert-Dussardier, L. Wittler, M. Borschiwer, S.A. Haas, M. Osterwalder, M. Franke, B. Timmermann, J. Hecht, M. Spielmann, A. Visel, and S. Mundlos. 2015. Disruptions of topological chromatin domains cause pathogenic rewiring of gene-enhancer interactions. Cell. 161:1012–1025.

Mach, P., P.I. Kos, Y. Zhan, J. Cramard, S. Gaudin, J. Tünnermann, E. Marchi, J. Eglinger, J. Zuin, M. Kryzhanovska, S. Smallwood, L. Gelman, G. Roth, E.P. Nora, G. Tiana, and L. Giorgetti. 2022. Cohesin and CTCF control the dynamics of chromosome folding. Nature genetics. 54:1907–1918.

Marshall, W.F., A. Straight, J.F. Marko, J. Swedlow, A. Dernburg, A. Belmont, A.W. Murray, D.A. Agard, and J.W. Sedat. 1997. Interphase chromosomes undergo constrained diffusional motion in living cells. Curr Biol. 7:930–939.

Miné-Hattab, J., and R. Rothstein. 2013. DNA in motion during double-strand break repair. Trends Cell Biol. 23:529–536.

Mizuguchi, T., G. Fudenberg, S. Mehta, J.M. Belton, N. Taneja, H.D. Folco, P. FitzGerald, J. Dekker, L. Mirny, J. Barrowman, and S.I. Grewal. 2014. Cohesin-dependent globules and heterochromatin shape 3D genome architecture in S. pombe. Nature. 516:432–435.

Moreno, S., A. Klar, and P. Nurse. 1991. Molecular genetic analysis of fission yeast Schizosaccharomyces pombe. Methods Enzymol. 194:795–823.

Munoz, S., M. Minamino, C.S. Casas-Delucchi, H. Patel, and F. Uhlmann. 2019. A Role for Chromatin Remodeling in Cohesin Loading onto Chromosomes. Mol Cell. 74:664–673 e665.

Neumann, F.R., V. Dion, L.R. Gehlen, M. Tsai-Pflugfelder, R. Schmid, A. Taddei, and S.M. Gasser. 2012. Targeted INO80 enhances subnuclear chromatin movement and ectopic homologous recombination. Genes & development. 26:369–383.

Nora, E.P., B.R. Lajoie, E.G. Schulz, L. Giorgetti, I. Okamoto, N. Servant, T. Piolot, N.L. van Berkum, J. Meisig, J. Sedat, J. Gribnau, E. Barillot, N. Bluthgen, J. Dekker, and E. Heard. 2012. Spatial partitioning of the regulatory landscape of the X-inactivation centre. Nature. 485:381–385.

Nuebler, J., G. Fudenberg, M. Imakaev, N. Abdennur, and L.A. Mirny. 2018. Chromatin organization by an interplay of loop extrusion and compartmental segregation. Proc Natl Acad Sci U S A. 115:E6697–E6706.

Ocampo-Hafalla, M., S. Munoz, C.P. Samora, and F. Uhlmann. 2016. Evidence for cohesin sliding along budding yeast chromosomes. Open Biol. 6.

Oshidari, R., J. Strecker, D.K.C. Chung, K.J. Abraham, J.N.Y. Chan, C.J. Damaren, and K. Mekhail. 2018. Nuclear microtubule filaments mediate non-linear directional motion of chromatin and promote DNA repair. Nature communications. 9:2567.

Parelho, V., S. Hadjur, M. Spivakov, M. Leleu, S. Sauer, H.C. Gregson, A. Jarmuz, C. Canzonetta, Z. Webster, T. Nesterova, B.S. Cobb, K. Yokomori, N. Dillon, L. Aragon, A.G. Fisher, and M. Merkenschlager. 2008. Cohesins functionally associate with CTCF on mammalian chromosome arms. Cell. 132:422–433.

Pradhan, B., R. Barth, E. Kim, I.F. Davidson, B. Bauer, T. van Laar, W. Yang, J.K. Ryu, J. van der Torre, J.M. Peters, and C. Dekker. 2022. SMC complexes can traverse physical roadblocks bigger than their ring size. Cell reports. 41:111491.

Rao, S.S.P., S.C. Huang, B. Glenn St Hilaire, J.M. Engreitz, E.M. Perez, K.R. Kieffer-Kwon, A.L. Sanborn, S.E. Johnstone, G.D. Bascom, I.D. Bochkov, X. Huang, M.S. Shamim, J. Shin, D. Turner, Z. Ye, A.D. Omer, J.T. Robinson, T. Schlick, B.E. Bernstein, R. Casellas, E.S. Lander, and E.L. Aiden. 2017. Cohesin Loss Eliminates All Loop Domains. Cell. 171:305–320 e324.

Schmidt, C.K., N. Brookes, and F. Uhlmann. 2009. Conserved features of cohesin binding along fission yeast chromosomes. Genome Biol. 10:R52.

Schrank, B.R., T. Aparicio, Y. Li, W. Chang, B.T. Chait, G.G. Gundersen, M.E. Gottesman, and J. Gautier. 2018. Nuclear ARP2/3 drives DNA break clustering for homology-directed repair. Nature. 559:61–66.

Seeber, A., V. Dion, and S.M. Gasser. 2013. Checkpoint kinases and the INO80 nucleosome remodeling complex enhance global chromatin mobility in response to DNA damage. Genes & development. 27:1999–2008.

Sexton, T., E. Yaffe, E. Kenigsberg, F. Bantignies, B. Leblanc, M. Hoichman, H. Parrinello, A. Tanay, and G. Cavalli. 2012. Three-dimensional folding and functional organization principles of the Drosophila genome. Cell. 148:458–472.

Shen, Y., F. Yue, D.F. McCleary, Z. Ye, L. Edsall, S. Kuan, U. Wagner, J. Dixon, L. Lee, V.V. Lobanenkov, and B. Ren. 2012. A map of the cis-regulatory sequences in the mouse genome. Nature. 488:116–120.

Stigler, J., G.O. Camdere, D.E. Koshland, and E.C. Greene. 2016. Single-Molecule Imaging Reveals a Collapsed Conformational State for DNA-Bound Cohesin. Cell reports. 15:988–998.

Swartz, R.K., E.C. Rodriguez, and M.C. King. 2014. A role for nuclear envelope-bridging complexes in homology-directed repair. Molecular biology of the cell. 25:2461–2471.

Symmons, O., V.V. Uslu, T. Tsujimura, S. Ruf, S. Nassari, W. Schwarzer, L. Ettwiller, and F. Spitz. 2014. Functional and topological characteristics of mammalian regulatory domains. Genome research. 24:390–400.

Terakawa, T., S. Bisht, J.M. Eeftens, C. Dekker, C.H. Haering, and E.C. Greene. 2017. The condensin complex is a mechanochemical motor that translocates along DNA. *Science (New York*, N.Y.).

Thon, G., P. Bjerling, C.M. Bünner, and J. Verhein-Hansen. 2002. Expression-state boundaries in the mating-type region of fission yeast. Genetics. 161:611–622.

Tiana, G., and L. Giorgetti. 2018. Integrating experiment, theory and simulation to determine the structure and dynamics of mammalian chromosomes. Curr Opin Struct Biol. 49:11–17.

Tran, P.T., L. Marsh, V. Doye, S. Inoué, and F. Chang. 2001. A mechanism for nuclear positioning in fission yeast based on microtubule pushing. J Cell Biol. 153:397–411.

Van Bortle, K., and V.G. Corces. 2012. tDNA insulators and the emerging role of TFIIIC in genome organization. Transcription. 3:277–284.

Wang, S., J.H. Su, B.J. Beliveau, B. Bintu, J.R. Moffitt, C.T. Wu, and X. Zhuang. 2016. Spatial organization of chromatin domains and compartments in single chromosomes. *Science (New York*, N.Y.). 353:598–602.

Weber, S.C., A.J. Spakowitz, and J.A. Theriot. 2012. Nonthermal ATP-dependent fluctuations contribute to the in vivo motion of chromosomal loci. Proc Natl Acad Sci U S A. 109:7338–7343.

Wendt, K.S., K. Yoshida, T. Itoh, M. Bando, B. Koch, E. Schirghuber, S. Tsutsumi, G. Nagae, K. Ishihara, T. Mishiro, K. Yahata, F. Imamoto, H. Aburatani, M. Nakao, N. Imamoto, K. Maeshima, K. Shirahige, and J.M. Peters. 2008. Cohesin mediates transcriptional insulation by CCCTC-binding factor. Nature. 451:796–801.

Zhang, X.R., L. Zhao, F. Suo, Y. Gao, Q. Wu, X. Qi, and L.L. Du. 2022. An improved auxin-inducible degron system for fission yeast. G3 (*Bethesda*). 12.

Zhurinsky, J., S. Salas-Pino, A.B. Iglesias-Romero, A. Torres-Mendez, B. Knapp, I. Flor-Parra, J. Wang, K. Bao, S. Jia, F. Chang, and R.R. Daga. 2019. Effects of the microtubule nucleator Mto1 on chromosomal movement, DNA repair, and sister chromatid cohesion in fission yeast. Molecular biology of the cell. 30:2695–2708.

Zimmer, C., and E. Fabre. 2019. Chromatin mobility upon DNA damage: state of the art and remaining questions. Current genetics. 65:1–9.

